# MINFLUX fluorescence nanoscopy in biological tissue

**DOI:** 10.1101/2024.07.06.602333

**Authors:** Thea Moosmayer, Kamila A. Kiszka, Volker Westphal, Jasmin K. Pape, Marcel Leutenegger, Heinz Steffens, Seth G. N. Grant, Steffen J. Sahl, Stefan W. Hell

**Affiliations:** Department of NanoBiophotonics, Max Planck Institute for Multidisciplinary Sciences, Göttingen 37077, Germany; Georg-August University School of Science (GAUSS), University of Göttingen, Göttingen, Germany; Centre for Clinical Brain Sciences, The University of Edinburgh, Edinburgh EH16 4SB, United Kingdom; Department of Optical Nanoscopy, Max Planck Institute for Medical Research, Heidelberg 69120, Germany

## Abstract

Optical imaging access to nanometer-level protein distributions in intact tissue is a highly sought-after goal, as it would provide visualization in physiologically relevant contexts. Under the unfavorable signal-to-background conditions of increased absorption and scattering of the excitation and fluorescence light in the complex tissue medium, super-resolution fluorescence microscopy methods are severely challenged in attaining precise localization of molecules. We reasoned that the typical use of a confocal detection pinhole in MINFLUX nanoscopy, suppressing background and providing optical sectioning, should facilitate the detection and resolution of single fluorophores even amid scattering and optically challenging tissue environments. Here, we investigated the performance of MINFLUX imaging for different synaptic targets and fluorescent labels in tissue sections of the mouse brain. Single fluorophores were localized with a precision of *<* 5 nm at up to 80 µm sample depth. MINFLUX imaging in two color channels allowed to probe PSD95 localization relative to the spine head morphology, while also visualizing presynaptic VGlut clustering and AMPA receptor clustering at the post-synapse. Our two-dimensional (2D) and three-dimensional (3D) two-color MINFLUX results in tissue, with *<* 10 nm 3D fluorophore localization, open up new avenues to investigate protein distributions on the single-synapse level in fixed and living brain slices.

---

The breaking of the diffraction resolution barrier (1) in fluorescence microscopy is an important development in the microscopy of cells and tissues. With fluorophore transitions between ‘on’ and ‘off’ states of fluorescence as the central element for feature separation (2), nanoscopy or super-resolution (SR) approaches have come in two distinct categories: STED-like (3, 4) coordinate-targeted and PALM/STORM-like (5, 6) coordinate-stochastic SR variants, each with specific strengths and weaknesses.

Aiming to extend the SR capabilities to a range of physiologically relevant contexts, confocal scanning STED imaging has been shown to be applicable deeper down in cells and tissues (7-9), but remains difficult to advance to highest (*<* 20 nm) resolution performance. The wide-field single-molecule based PALM/STORM methods reach such resolution somewhat more readily, but face challenges inside tissues and at larger optical depth. Total-internal reflection (TIRF) conditions (close to the coverslip surface), and/or other background-mitigating strategies such as selective plane illumination are effectively required for thicker samples, including tissue applications. In combination with optical sectioning, PALM/STORM was reported to yield 50−70 nm resolutions in up to 15 µm imaging depth within living cells and organotypic tissues by employing oil immersion and “digital pinholing” (10), or in 40-60 µm thick fixed tissue slices when utilizing a water immersion lens and angled light-sheet illumination (11). Fluorophore localization by finding the center of the diffraction spot of the fluorescence of individual fluorophores on a camera, as fundamentally used in PALM/STORM, is compromised by background when the fluorophores are located deep inside tissue.

MINFLUX nanoscopy combines the strengths of the coordinate-targeted and -stochastic approaches. While the fluorescence capability of individual fluorophores is stochastically turned on and off, MINFLUX probes the position of a fluorophore in the sample with a focal excitation pattern containing an intensity minimum, such as a doughnut-shaped focus. The closer the minimum is to the fluorophore, the fewer fluorescence photons are needed to localize the fluorophore. In a way, the burden of requiring many photons for precise optical localization is shifted from the feeble stream of fluorescence photons to the stable and bright beam of the laser. This fundamental trait of MINFLUX localization brings about major advantages as much fewer fluorescence photons are needed to localize a fluorophore with a certain precision. Only the intensity profile of the excitation pattern around the minimum needs to be sufficiently maintained for precise localization, with much less stringent requirements on the detection path. Last but not least, the typical use of a confocal detection pinhole in MINFLUX suppresses background and provides some optical sectioning, facilitating the detection of single fluorophores amid a scattering environment such as deeper regions of tissue. Therefore, unlike in PALM/STORM, localization of single fluorophores should be possible deeper down in scattering media. Hence, we decided to investigate if and to which extent MINFLUX is able to image at the nanoscale deeper inside tissue.

Other approaches to achieve nanoscopy in tissue were reported, such as expansion microscopy (12). As they require harsh chemical alterations of the sample as part of the workflow, their development would never lead to live-cell imaging. In contrast, MINFLUX deserves development because it leaves the sample morphologically largely intact and is inherently compatible with live-cell imaging. Here, we take the first step to explore this direction. We pioneer MINFLUX imaging in tissue and show that it is possible to obtain resolution, i.e., detail clarity, at the nanometer scale. A major step towards live-tissue nanoscopy, our experiments clarify and address relevant practical challenges and pave the way for its optical feasibility.

Specifically, we describe MINFLUX imaging in tissue sections from mouse brain, featuring *<* 5 nm localization precisions at up to 80 µm depth, by showing fluorescently labeled caveolin-1 distributions, and by visualizing actin and PSD95 at the post synapse at several tens-of-micrometer depth. The potential for live-tissue imaging is exemplified in experiments with PSD95, and MINFLUX in two color channels allowed to probe PSD95 localization relative to the morphology of spine heads. MINFLUX visualizes presynaptic vesicular glutamate transporter (VGlut) clustering and the clustering of α-amino-3-hydroxy-5-methyl-4-isoxazolepropionic acid receptors (AMPAR) at the post-synapse. Extensions to 3D imaging of actin, PSD95 and AMPA receptors demonstrate the potential for investigations of proteins in the volume and at the surface of dendritic spines with remarkable 3D precisions of ∼ 5 nm laterally (*xy*) and 10-15 nm axially (*z*) in the complex tissue setting.

## Results

### Adaptations of experimental setup

To date, MINFLUX imaging (13-17) has been demonstrated in relatively flat cells cultured in monolayers, or single immobilized retina layers directly on the coverslip (18). Imaging deeper in tissue comes with a decreased signal-to-background ratio (SBR), because of signal loss due to absorption and scattering in the tissue, an increased background from out-of-focus fluorophores and contributions from tissue auto-fluorescence. Additionally, for highly precise localizations, the sample position needs to be stabilized against drifts in all imaging depths of interest, which is not possible with the MINFLUX focus lock systems described previously (13, 15). Furthermore, the refractive index mismatch between tissue and immersion oil leads to spherical aberrations, which pose challenges to the control of the focal intensity distribution and the clean formation of its minimum, especially for 3D MINFLUX.

The elements of our custom-built beam-scanning MINFLUX system are shown in Fig.1A and more detailed in Fig. S1 (*Supplementary Information*). Aside from a number of key components required for efficient operation of any MINFLUX nanoscope, such as suitable laser sources, fast beam deflection devices for fluorophore targeting, a fluorescence scanner and dedicated real-time hard- and software control, several features make our modified implementation more suitable for tissue imaging. Importantly, we chose to use a silicone oil-immersion objective for improved refractive index matching and aberration compensation (compare Supplementary Text, *Supplementary Information*).

**Fig. 1.**
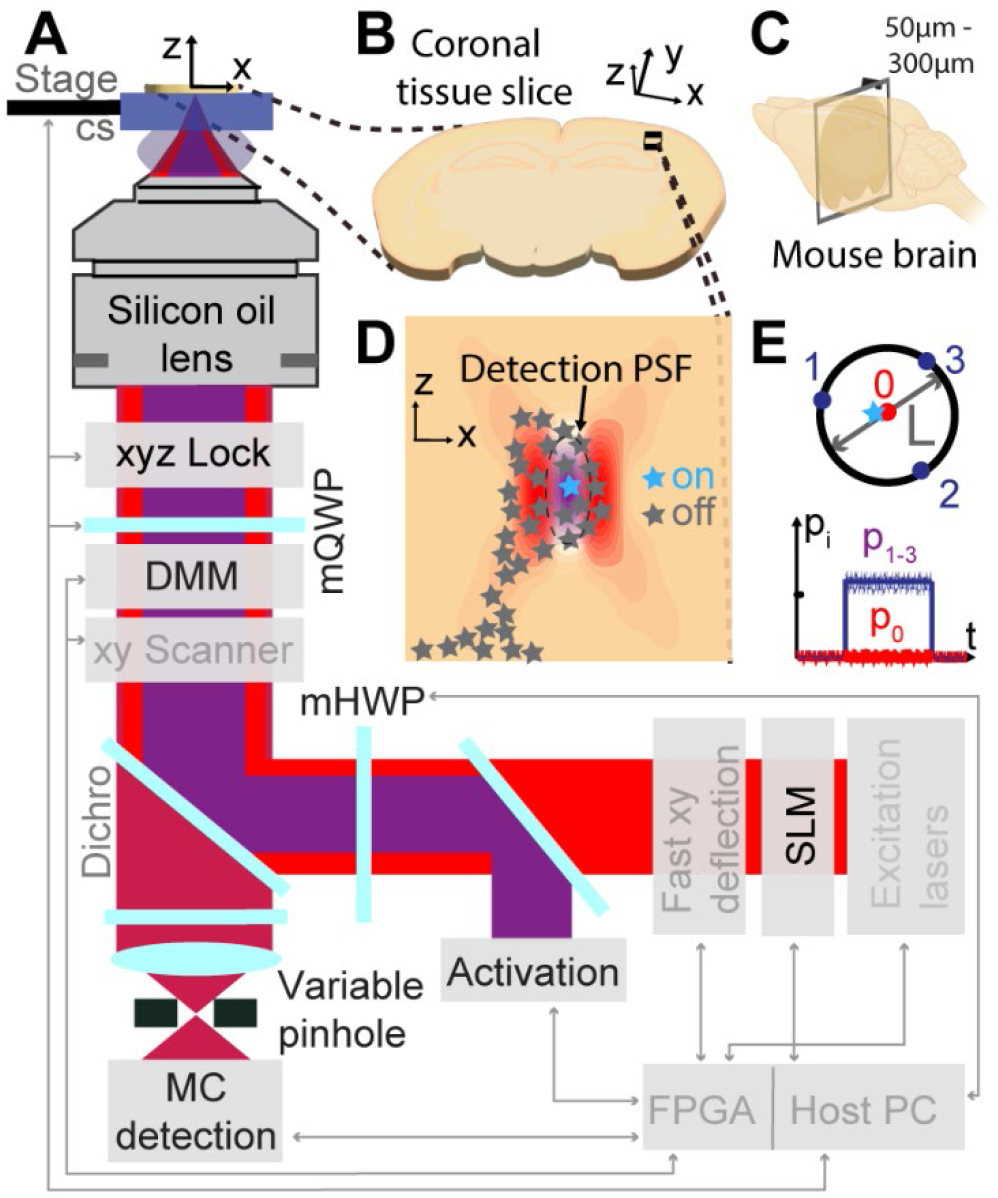
MINFLUX nanoscopy in biological tissue. **A** Overview of the custom-built MINFLUX setup for imaging in tissue. Adaptations for tissue imaging are highlighted in black, standard MINFLUX components shown in gray font. **B** Sample imaged. The tissue is taken as coronal section from a mouse brain as depicted in **C. D** Region of interest in the brain slice showing the optical sectioning capabilities of MINFLUX by the confocal detection volume and the selective on- and off-switching of single emitters. The activated fluorophore is localized photon-efficiently by centering the zero of the doughnut excitation beam on the activated molecule. The targeted-coordinate pattern for the doughnut excitation beam and the schematic photon emission trace of a centered molecule are shown in **E**. cs – coverslip, PSF – point spread function, DMM – deformable membrane mirror, mQWP – motorized quarter-wave plate, mHWP – motorized half-wave plate, p_i_ – photon count ratio in i^th^ exposure, MC – multicolor, FPGA – field-programmable gate array, SLM – spatial light modulator.

The instrument employs a confocal pinhole with variable size, allowing to adapt to samples of varying fluorophore density and to realize suitable SBR for single-fluorophore detection (Fig. S7). Motorized achromatic quarter- and half-wave plates (QWP/HWP) control the polarization of the excitation light and enable MINFLUX localizations with a flexible set of different excitation wavelengths (511 nm, 560 nm and 647 nm) in combination with the photodetectors in a multicolor detection unit.

A deformable membrane mirror (DMM) as part of the xyz-scanning system allows for fast focus shifts and therefore efficient 3D MINFLUX localizations. The DMM also enables fast aberration correction of the excitation and detection point spread function (PSF) simultaneously. A spatial light modulator (SLM) enables aberration correction of the excitation PSF. There is also an option to provide oxygen for living tissue slices (Fig. S1). Since stability of the sample with respect to the objective lens is critical, we developed an active depth-adaptable focus lock system. Furthermore, we implemented a progressive activation scheme that automizes and speeds up the measurement process. The depth-adaptable focus-lock system and the progressive activation are described in detail in the next section.

Samples were usually coronal slices of mouse brain tissue with 60-300 µm thickness (Fig. 1B). A cut through the brain leading to a coronal section is indicated in Fig. 1C. An imaging region of interest in the brain tissue section can be selected. We chose imaging regions in the cortex or hippocampus. This region of interest was then scanned using a piezo-driven tip-tilt mirror, and sparse fluorophores in the ‘on’ state (Fig. 1D) were localized by centering the excitation doughnut onto them. The centering (MINFLUX localization) was accomplished by iteratively moving the targeted-coordinate pattern (TCP) onto the fluorophore of interest while shrinking the geometric parameter *L* that defines the diameter of the TCP (Fig. 1E). In each MINFLUX iteration, the multiplexed emission trace provided a more precise fluorophore coordinate (Fig. 1E). If a fluorophore is successfully centered in the TCP, the ratio p_0_ of the number of detected photons n_0_ in the central doughnut exposure divided by the number of photons *N* collected in all four exposures, should be (close to) zero (compare Fig. 1E). All details of the utilized MINFLUX localization sequences can be found in Tables S3-S7 (*Supplementary Information*).

Both the selective photoactivation of individual emitters in the brain tissue slice and the confocal detection (Fig. 1D) contribute to optical sectioning by reducing the out-of-focus background. Selective photoactivation also reduces the bleaching of out-of-focus fluorophores. Filtering of the photon emission traces for emission events during which the fluorophore of interest was centered provides a further means to discard background (Material and Methods, and Table S8, *Supplementary Information*).

### Depth-adaptable focus lock system and progressive activation

To allow nanometer-precise localizations, the MINFLUX setup needs to be stable. This concerns mechanical components (vibrations, thermal drift, stage settling), beam positions (beam jitter due to air turbulence and sound) as well as beam power. The need to actively move the sample with the stage to find the region of interest (ROI) for a measurement typically resulted in the sample stage and sample not being equilibrated and, consequently, a main source of drift during the MINFLUX measurement, which lasted typically tens of minutes for the ROIs in this work. To reduce this drift, active drift correction of the sample with respect to the objective has been commonly employed in MINFLUX imaging (13, 15). The strategy is monitoring and controlling the position of fiducial markers on the coverslip or directly the coverslip and therefore only works for imaging in close proximity to the coverslip.

We constructed a focus lock system that can be adapted to different imaging depths, consisting of two independent parts for stabilizing the sample position perpendicular to the optical axis (the *xy*-lock system, Fig. 2A) and for stabilizing the sample position along the optical axis (the *z*-lock system, Fig. 2B). For *xy*-position stabilization (Fig. 2A), the light from a near-infrared superluminescent LED (∼850 nm) was focused into the back aperture of the objective lens, producing a collimated beam to illuminate an area of about 50×50 µm^2^ on the sample coverslip. The gold nanorods applied to the coverslip scattered the light, effecting polarization changes depending on the orientation of the nanorods. While the back-reflection from the sample interfaces did not pass back over the D-shaped mirror and was not deflected by the polarizing beam splitter (PBS), the back-scattered light from the nanorods did return over the D-shaped mirror and was deflected by the PBS (if its polarization was changed due to nanorod orientation) and then imaged onto a camera by a lens with variable focal length. This variable focus allowed sharp imaging of the nanorods at different axial sample placements. Thereby, active sample stabilization at different imaging depths was achieved.

**Fig. 2.**
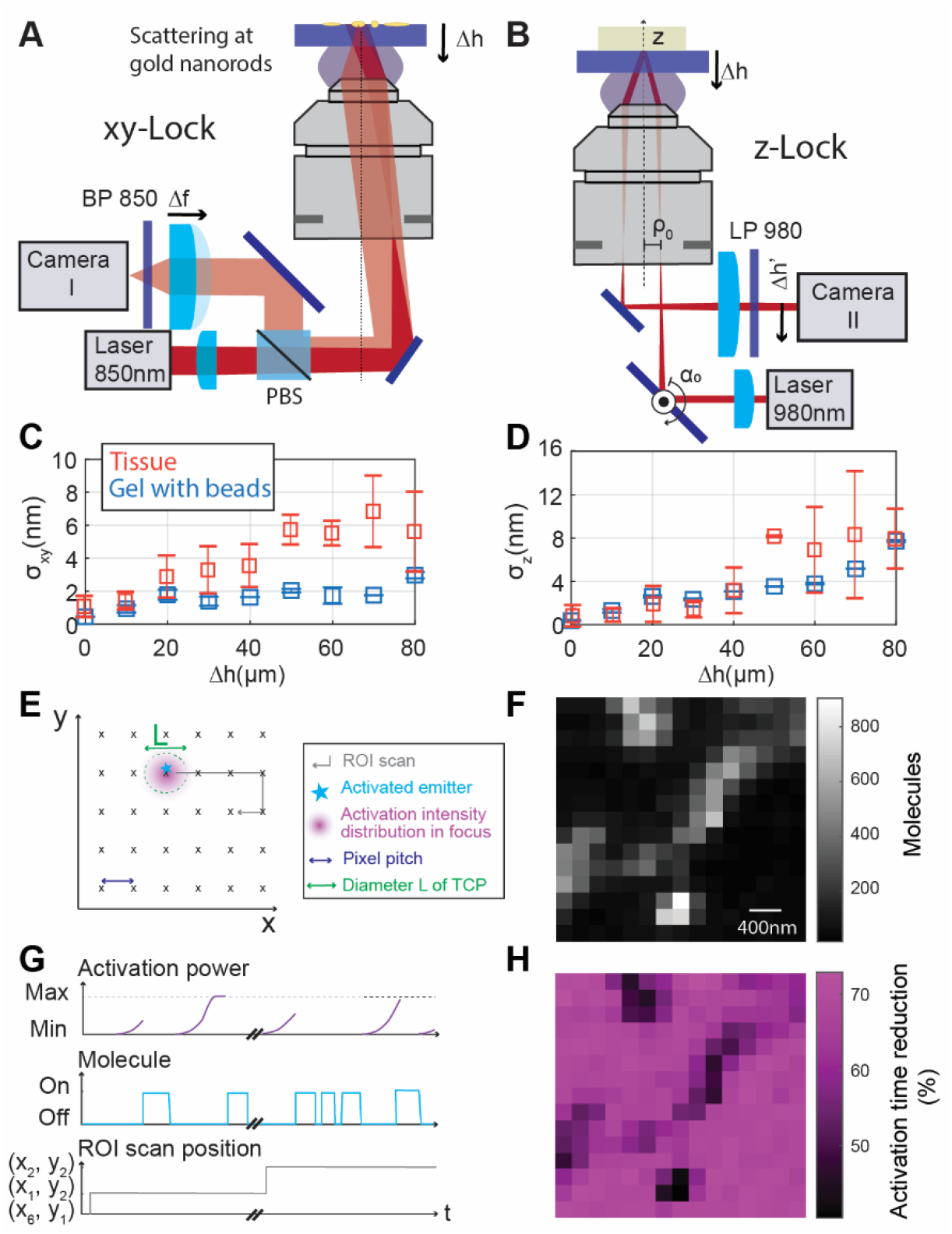
Active drift-correction for depth imaging and progressive activation. **A** Depth-adaptable xy-lock system. The dark-field images of fiducial markers (gold nanorods) on the coverslip can be acquired when imaging in a plane away from the coverslip surface by refocusing the nanorods with a variable-focus lens. **B** Depth-adaptable z-lock system. The imaging depth range that can be stabilized with high precision is enlarged by a piezo-actuated closed-loop absolute positioning mirror mount. (**C**,**D**) Stability of the **C** xy-lock position and **D** z-lock position for measurements in both agarose-sucrose samples and tissue samples. Mean stability values and standard deviation shown. **E** Schematic of activation scan over the ROI. **F** Progressive activation, shown schematically. At each new activation location addressed by the mosaic scan, the activation power is ramped up until a molecule starts to emit. **F** Number of fluorophores detected at each mosaic scan position. **G** Residual activation time needed for the progressive activation in percent of activation time expected with constant low activation. PBS – polarizing beam splitter, BP – band pass, LP – long pass, Δh – imaging depth, α_0_ – rotation angle of absolute positionable mirror, Δh’ – z-lock beam position on camera, σ_xy_ – xy-lock position stability, σ_z_ – z-lock position stability, max – maximum, min – minimum.

The position of selected nanorods in the live camera image was fitted by a Gaussian distribution and the position was stabilized by an active feedback loop with a highly precise (0.1 nm positioning resolution) piezo stage (P-733.3DD, Physik Instrumente). The position stability of the xy-lock system was characterized by quantifying the x- and y-deviations from the mean over time (Fig. S3 A,B).

The stabilization performance was investigated for measurements in different imaging depths and in different samples. The lock precision in depth in an agarose-sucrose gel (*n* = 1.38; matching the refractive index of mouse brain tissue in layer 1 of the cortex) with embedded fluorescent beads is shown in Fig. 2C, along with measurements in different tissue samples for measurement durations of up to 2h. The focus lock stability (root-mean-square uncertainty) *σ*_*xy*_was bounded to < 2 nm in the gel sample and ≲ 6 nm in the tissue at up to 80 µm depth. In practice, fluorophore localization uncertainties below 5 nm could be observed for short measurement times in small fields of view (i.e. caveolin samples, see below).

The stability of the focus lock at optical depth was independently cross-checked by monitoring the position of fluorescent beads or labeled structures via the main imaging system. Of note, the stability was lower in the tissue sample than in the gel sample, mainly due to scattering of the incoming light in the tissue leading to a reduced SBR on the camera. The tissue is also less homogeneous than the gel sample, leading to a larger spread in the stability among measurements.

For stabilization along the optical axis (*z*), a separate NIR (980 nm) laser beam was focused into the back aperture of the objective lens and impinged collimated on the coverslip-sample interface obliquely (Fig. 2B). The reflected light from the coverslip-sample interface passed collimated onto a camera. Using the ray transfer matrix analysis and approximating the objective lens by a thin lens, the position Δ*h*^′^of the reflected light on the camera was calculated in a simplified way from the starting parameters ρ_0_ (displacement from optical axis) and α_0_ (angle of incidence) of the incident beam before the objective back aperture, yielding

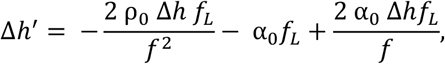

with the imaging depth Δ*h*, focal length *f* of the objective lens and the focal length *f*_*L*_ of the lens in front of the camera. For all other parameters kept constant, a movement of the sample along the optical axis directly translated into a movement of the reflected beam on the camera. This effect was used for the active stabilization. A proportional-and-integral (PI-) control loop moving the piezo fine stage actively stabilized the center position of the Gaussian intensity distribution on the camera.

There is a trade-off between sensitivity and depth range of the z-lock system, because the maximal range of beam movement Δ*h*’ was limited by the camera chip size (∼4.5 mm). By changing the angle of incidence α_0_ by a piezo mirror, we were able to reliably and precisely switch between a regime with high *z* sensitivity and a regime with larger depth range (Fig. S3C). In subsequent measurements, we routinely adjusted the system to the high-sensitivity regime for 0-15 µm depth and to a lower-sensitivity regime for 15-80 µm depth.

When testing the focus lock system in the *n* = 1.38 gel sample, we documented a slight deterioration of the position stability σ_*z*_ along the optical axis when focusing deeper into the sample, at < 4 nm until 60 µm depth and ∼8 nm at 80 µm depth (Fig. 2D). This was observed to be due to a lower reflectivity of the immersion-oil-to-sample interface when adjusting the *z*-lock for the depth, leading to lower SBR and higher uncertainty of the position. When adjusting α_0_ to the depth, this also led to a slight defocusing of the beam, further decreasing the SBR. We also measured the focus lock precision in tissue samples, observing a similar trend (Fig. 2D). At large depth, we noticed lower stability, probably due to reduced SBR caused by the light scattering and absorption of the tissue.

We observed that, when calibrated, and for coverslips of known thickness, the beam position in the z-lock system was another means to determine and confirm the imaging depth in the sample besides the position sensor from the coarse and fine stages.

To speed up the MINFLUX acquisitions and to adapt the activation power to inhomogeneous fluorophore densities (Fig. 2F), we implemented a progressive activation scheme (Fig. 2E, G). The activation power of the 405 nm laser – which served to re-activate both blinking organic fluorophores and photoconvert fluorescent proteins – was ramped up exponentially following each detected fluorophore and at each new activation location in the sample. With a low starting activation power, the likelihood of several fluorophores being activated at the same time was reduced while maintaining a high fluorophore detection rate due to the about 100-fold ramped-up activation. As the progressive activation procedure acts locally during scanning in a sample-responsive manner, it inherently reduced the time expended in locations where no new fluorophores were activated, whereas it exhausted the fluorophores in crowded locations by activating them successively. To calculate the reduction in activation time due to progressive activation (Fig. 2F and *Supplementary Information*, Fig. S11), the time needed to activate a molecule with progressive activation was compared to the estimated time needed with constant low activation to reach the same activation dose. Typically, the reduced activation times led to 50-75% shorter image acquisition durations.

### MINFLUX nanoscopy up to 80 µm deep in tissue

To examine performance trends in 2D MINFLUX tissue imaging, we imaged caveolin-1 labeled with primary and secondary antibody conjugated to the dye Alexa Fluor 647. A well-known blinking buffer for this established dye system was used. Caveolin-1 is a membrane protein found in most cells. It is linked to the formation of so-called lipid rafts and the formation of caveolae, which are small (50-100 nm diameter), often lightbulb-shaped invaginations of the cell membrane or of spherical vesicles in the cell (19-21). Although neurons are one of the few cell types that do not contain caveolae structures, the Caveolin-1 protein is found in neurons, where it is linked to the plasticity of neurons and especially synapses (22).

To investigate differences between MINFLUX imaging in flat cells and complex tissue, we compared Caveolin-1 images and MINFLUX metrics in U2-OS cells and tissue slices directly on the coverslip (Fig S5). The median SBR was considerably reduced in tissue (factor >2). Furthermore, the left peak in the histogram of the p_0-_distribution shifted to the right.

We independently checked the emission trace during MINFLUX localization of isolated, single molecules to quantify and separate different influences on the p_0_ distribution: (i) The light intensity in the central minimum of the doughnut is not strictly zero, but some residual excitation light remains (at least ∼1 % of single-molecule emission is elicited compared to the maximum at the donut crest). (ii) There is background from the system, such as dark counts from the avalanche photodiode (APD) detectors, auto-fluorescence and scattering from the sample, which can be nearly completely suppressed by spectral filtering (a contribution of ∼0.7 % was determined by single-molecule PSF measurements of the ‘zero quality’ with stepwise bleaching of single fluorophores to the background level; measurement at the glass surface). (iii) There is fluorescence from other fluorophores which activate spontaneously outside the targeted volume or do not switch off fully, and whose fluorescence contribution can be only partly suppressed by the confocal detection. This contribution from other fluorophores is deemed to be the most prominent source of background in MINFLUX localizations in a real cell or tissue sample.

With increasing depth Δh from the coverslip, the focal doughnut pattern became more elongated in the axial direction, and the intensity contrast between doughnut crest and minimum was reduced, from ∼3 % at the glass surface to ∼14 % residual intensity at the center of the pattern at 80 µm depth (Fig. 3A insets in the respective left panels; constant background offset in the image subtracted). A more detailed overview of focal-region doughnut shapes in different sample depths can be found in Fig. S4 and the accompanying SI Supplementary Text.

**Fig. 3.**
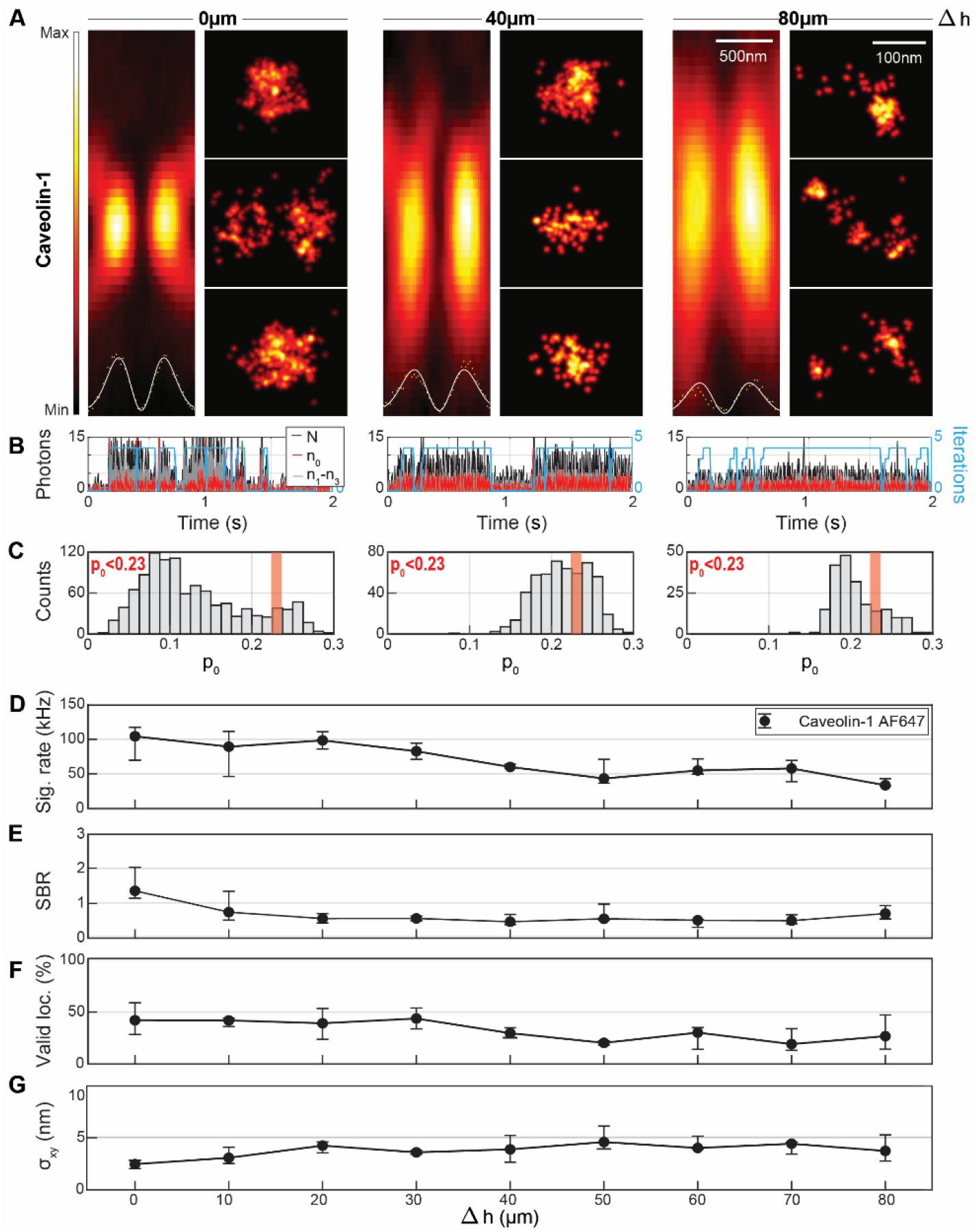
Imaging up to 80 µm deep in tissue with MINFLUX nanoscopy. **A** Focal intensity distribution (xz) of the excitation doughnut in different imaging depths in tissue together with regions of interest selected from MINFLUX images in the same imaging depth, showing caveolin-1 distributions. Intensity profiles of the excitation light are shown as insets together with a doughnut-shaped fit of the intensity along the x-axis. Intensity minimum deteriorates from 3 % at the coverslip to 14 % in 80 µm depth. The FWHM of the focus along *z* increases from ∼900 nm at the coverslip to ∼1660 nm in 80 µm depth. **B** Photon traces are shown for the upper row of MINFLUX images. **C** Histogram of *p*_*0*_ values for resegmented localizations. **D** Median signal count rate and interquartile range in each imaging depth. **E** Median SBR over depth. **F** Median ratio of valid localizations over depth. **G** Median localization precision over depth. Statistics calculated from 703924 localizations (23 images) in ∼0 µm, 58303 localizations (9 images) in ∼10 µm, 50396 localizations (10 images) in ∼20 µm, 10051 localizations (2 images) in ∼30 µm, 16009 localizations (5 images) in ∼40 µm, 53764 localizations (5 images) in ∼50 µm, 32978 localizations (6 images) in 60 µm, 193465 localizations (15 images) in 70 µm and 12289 localizations (7 images) in ∼80 µm imaging depth. Δh – imaging depth, N - number of photons collected in one multiplexing cycle. n_i_ – number of photons collected in i^th^ exposure, p_0_ = n_0_/N, sig. – signal, loc. – localizations, σ_xy_ – localization precision.

The MINFLUX reconstructions were found to be comparable to those in previous reports showing fluorescence nanoscopy of caveolae in cells (23, 24) (Fig. 3A), similar in morphological appearance, overall dimensions and the numbers of contained localizations. Single-fluorophore signals above the background level (Fig. 3B) showed a decreasing trend with depth (see below), and the histogram of *p*_*0*_ values (Fig. 3C) exhibited a shift to higher *p*_*0*_ with depth, still yielding robust numbers of localizations below the background threshold to enable meaningful reconstructions. From Δh = 0 to 80 µm, with increasing depth, the average signal rate decreases from ∼100 kHz to ∼40 kHz (Fig. 3D). The attained SBR, while between 1 and 2 at the surface, was found to plateau at ∼0.5 beyond 20 µm depth (Fig. 3E), still allowing robust localizations. The ratio of valid localizations according to a *p*_*0*_ < 0.23 background filtering criterion decreased by up to 25% over the examined depth range, reaching a plateau at ∼25% (Fig. 3F). The results reveal that 40 or 80 µm deep in tissue the background level indeed is higher than at the coverslip surface, where there are only fluorophores in the light cone above the focus position that can contribute to the background, and the signal decreases due to increased absorption and scattering of the excitation and fluorescence by the tissue.

These effects caused a decline in the SBR and the shift of the signal peak in the *p*_*0*_ distribution closer to the background peak, making a clear separation based on *p*_*0*_ less straightforward with increasing depth. Aberrations introduced by local inhomogeneities in the refractive index and the refractive index mismatch between tissue and immersion oil – even for the more suitable silicone oil system that we utilized – further shifted the signal peak in the *p*_*0*_ distribution to the right. The median localization precision decreased with depth, from ∼2.5 nm to still < 5 nm at Δh = 80 µm (Fig. 3G), as was expected for a trend to lower SBR. Robustly attainable precision below 5 nm at many tens of micrometer depth appears remarkable and promising for further investigations. As demonstrated above, part of this experimentally determined precision can be attributed to imperfect lateral stabilization, as the measured SBR would still allow for a better localization precision (compare Fig. S6).

### MINFLUX nanoscopy of post-synaptic proteins in fixed tissue

The most important actor during formation of the post-synaptic terminal is actin, which is employed in the build-up and motility of the post-synapse (25). There are several receptors on the post-synapse that modulate signal transmission. These are held in place and arranged by the post-synaptic density, with major component PSD95 acting as a scaffold. PSD95 is also linked to synapse formation (26), and has been found to redistribute along the dendrite and migrate to synapses (27). We further evaluated the 2D MINFLUX performance for imaging actin and PSD95 in the mouse brain tissue at different imaging depths (Fig. 4).

**Fig. 4.**
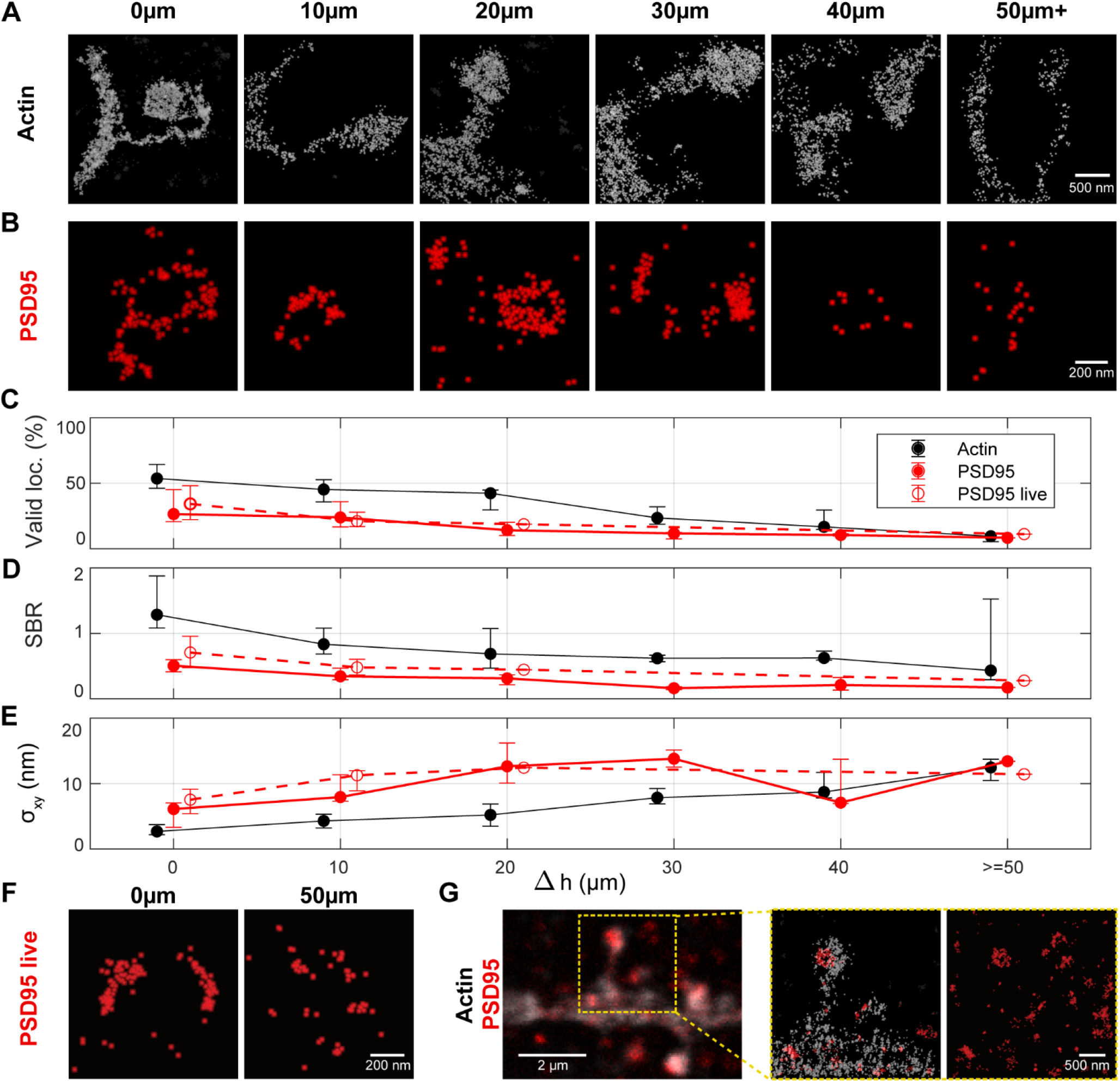
Imaging the post synapse deep in tissue. **A** MINFLUX nanoscopy of actin (LifeAct-GFP labeled with primary and secondary antibody by AF647) in 0-50 µm imaging depth. Localizations classified by delineation as not belonging to spine or dendrite are grayed out. **B** PSD95-mEOS2 in fixed tissue in 0-50 µm imaging depth. **C** Median percentage of valid MINFLUX localizations (p_0_ < 0.23) and interquartile range over imaging depth, calculated from several images in each imaging depth. **D** Median signal to background ratio over imaging depth. **E** Median localization precision over imaging depth. Number of localizations and images from which the statistics for the three different samples are calculated are shown in Table S10. **F** PSD95-mEOS2 in living tissue in 0 and 80 µm imaging depth. **G** Imaging with two excitation colors. Two-color image of PSD95-mEOS2 and actin-AlexaFluor647. From left to right: Two color confocal overview image. Two excitation color MINFLUX in the region of interest showing the PSD95 distribution on the post-synapse (only delineated dendritic region shown). Full image from the PSD95-mEOS2 color channel. σ – localization precision, SBR – signal to background ratio, loc. – localizations.

In a subset of neuronal dendrites, actin was labeled by viral injection of LifeAct-EYFP, which was then recognized by indirect immunofluorescence post-fixation with Alexa Fluor 647 as the fluorescent dye. LifeAct, labeling G- and F-actin, yields a volumetric filling of the dendrites and spines. In practice, we were able to acquire confocal previews in the EYFP and the Alexa Fluor 647 channel prior to MINFLUX measurements, which allowed to ascertain the structure of interest with reasonable confidence and exclude spots of apparent unspecific labeling. Judging from the confocal images acquired, there was indeed unspecific labeling visible in the Alexa Fluor 647 channel (but absent in the corresponding regions for EYFP).

MINFLUX visualized these small structures of characteristic morphology based on the actin content (Fig. 4A), that can be roughly divided into a spine head (as part of the synapse) and a spine neck connecting the synapse to the dendrite. As a clear border of the spines in the MINFLUX images could be identified, and only actin inside the dendrites should be labeled, we delineated the spines and rejected outliers from unspecific antibody labeling or detached dyes for improved visualization of the data (Fig. S8, *Supplementary Information*).

To approach the aim of measuring quantitative nanometer-precise protein distributions in living tissue, we investigated endogenous labeling of a protein of interest with a fluorescent protein (FP) by direct (covalent) labeling of proteins. This approach, while utilizing a dimmer fluorescent emitter, avoids the rather large size of antibodies, is live-cell compatible and stoichiometrically labels the protein. We thus performed MINFLUX on PSD95 endogenously labeled by mEOS2 (28) in different imaging depths (Fig. 4B). With an FP directly fused to the PSD95 molecules, we no longer observed unspecific labeling. mEOS2 is a photo-convertible fluorescent protein. Before photo-conversion (in its green-emitting form), we excited it at 511 nm wavelength to find ROIs containing PSD95. Then the mEOS2 were photo-converted individually with the 405 nm laser light and subsequently excited at 560 nm in their orange-emitting form. We initially checked the photo-conversion behavior in confocal imaging for each new tissue slice.

In the MINFLUX acquisitions of PSD95-mEOS2 (parameters in Tables S3 and S4, *Supplementary Information*), the emission events of the FP were confirmed to be brighter and more stable in the presence of an appropriate buffer (a 50 mM TRIS buffer in deuterium (29, 30), or the aforementioned blinking buffer) compared to the emission events in standard buffers such as PBS or ACSF. On average, the FP mEOS2 delivered ∼19 % fewer fluorescent photons and a ∼81 % lower photon emission rate than Alexa Fluor 647 and exhibited different switching kinetics. Therefore, we used modified parameters for the post-processing of the collected photon traces. A comparison of the fluorophore-adapted parameters is shown in Table S4 (*Supplementary Information*).

Similar trends to the caveolin-1 labeled samples, at similar performance with depth, were observed for actin and the FP-labeled PSD95 regarding the fraction of valid localizations, SBR and attained localization precision (Figs. 4C-E). We note that, at ≲ 10 nm in up to 50 µm, the demonstrated localization precision was slightly worse for mEOS2 than for Alexa Fluor 647. This can be mainly explained by localizing with half as many (∼1000) photons (mEOS2) compared to the dye case, yielding 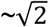 worse localization precision.

Compared to the earlier caveolin imaging, higher numbers of fluorophores in the ROI and therefore increasing MINFLUX measurement times for the sequential scanning posed additional high demands on the focus lock system that needs to be stable over the whole measurement time. As valid localization returns are reduced away from the glass and for increased background from a higher number of emitters out of focus, the time expended was further prolonged (to tens of minutes and up to 1-2 hours).

Taken together, the MINFLUX data of caveolae and dendritic spines show that 2D MINFLUX with single-digit nanometer localization precision is possible in several tens of micrometers depth. As shown for PSD95-mEOS2, this MINFLUX performance can also be obtained at the bluer wavelength and with a less bright emitter. A diversity of spine morphologies was resolved, with fine image detail resolvable (compare the neck of the left spine in Fig. 4A). PSD95 localization patterns in the imaged examples were of a high diversity, reflecting both a putative structural heterogeneity and likely planar projection (see 3D imaging below).

### Live tissue imaging

An FP genetically expressed in the mouse provides the possibility to image in living tissue. As we expected severe movement due to breathing and heart beat in the living mouse (7), we performed MINFLUX imaging on acute tissue slices. We kept the slices in ACSF buffer and provided oxygen during measurements to keep the slices alive for several hours. Imaging in several depths in living tissue slices gave comparable results as in fixed tissue slices (examples of two depths in Fig. 4F). An only slightly worse SBR and localization precision at the same imaging depths compared to fixed-slice imaging can be rationalized by a less optimal buffer system for the FP in acute slices, and by background from possibly not fully washed-out blood. Noticeably, the localization precision was slightly worse than in fixed slices even if the SBR was comparable or slightly more favorable. Since PSD95 is expected to show some movement *in vivo* (27, 31), this slightly worse localization precision was probably due to movement of the proteins. The measurement time for a MINFLUX image of PSD95 with a field of view of 3.2 × 3.2 µm^2^ was on average ∼20 minutes.

### MINFLUX nanoscopy with two excitation wavelengths

For imaging postsynaptic proteins in context, we established a robust multicolor imaging scheme, employing spectral separation both in detection (14) and excitation wavelength. The aforementioned fluorophores featured clearly separated absorption and emission spectra for this task and allowed sequential localization. An example of a two-color acquisition of actin and PSD95 is shown in Fig. 4G. The dendrite can be readily assigned and the corresponding PSD95 cluster selected.

Here, we localized first PSD95 and then actin. The PSD95-mEOS2 acquisition was therefore not affected by the actin imaging. Since both fluorophores can be activated by 405-nm light, the second image may be affected by the first imaging round. This effect was not expected to be very detrimental. As the blinking dye can be switched on multiple times, we should register the majority of dyes in any case. Note that a more conservative approach would be to first image Alexa Fluor 647, without photo-activation, until no more events are obtained, then image mEOS2, and then the dye again. We tested this approach for the 3D two-color imaging (see below).

### Analysis of 2D distribution of VGlut and AMPA receptors

Further molecular components of the synapse can be visualized. For illustration, we chose to image the arrangement of VGlut in synaptic vesicles of the pre-synapse (32) and AMPARs (33) on the post-synapse (Fig. 5A).

**Fig. 5.**
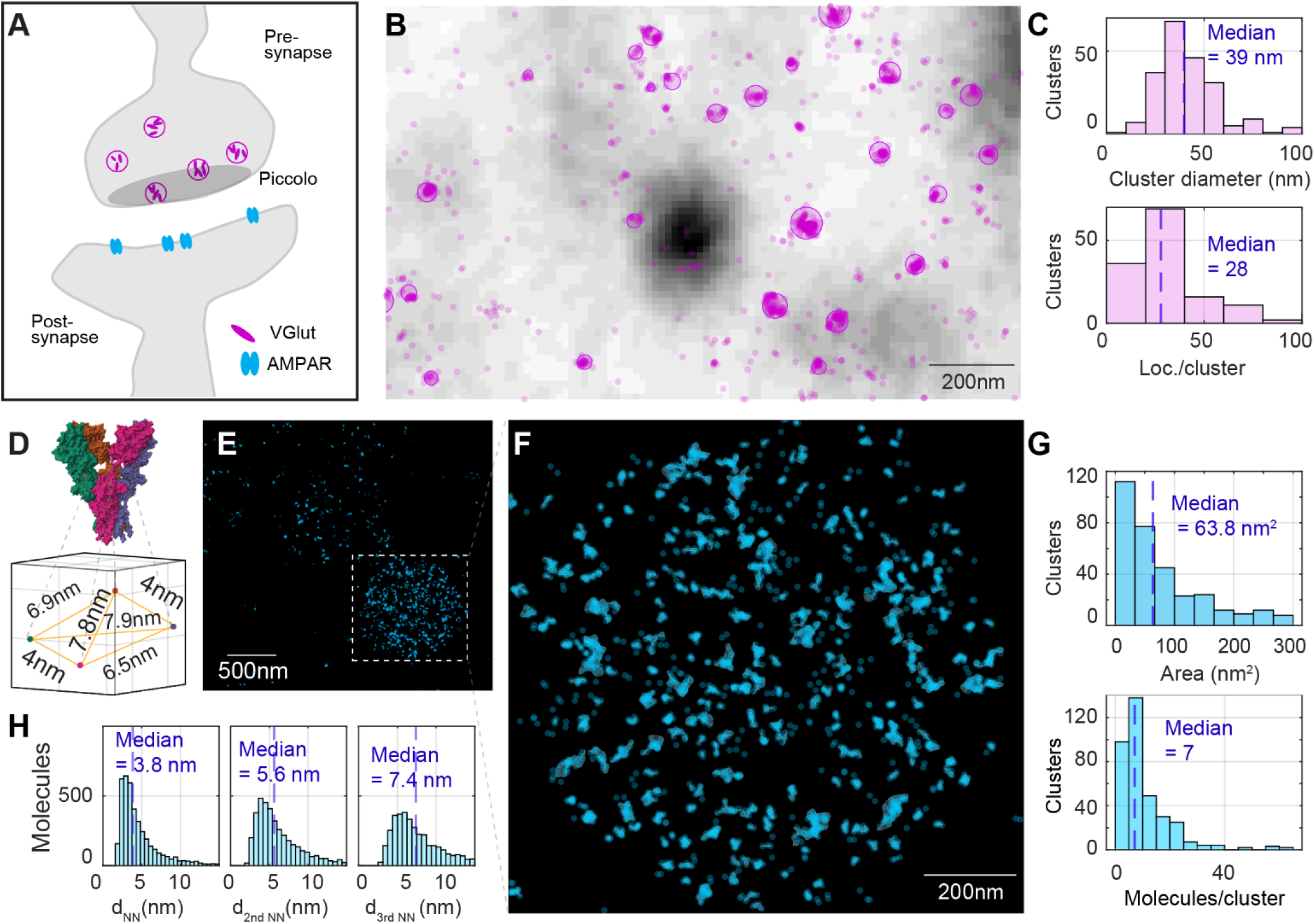
Cluster analysis of synaptic proteins in mouse brain tissue. **A** Schematic drawing of a synapse showing the arrangement of VGlut in synaptic vesicles of the pre-synapse, piccolo situated close to the synaptic release site, and AMPA receptors on the post synapse. **B** Confocal image of Piccolo (grayscale) overlayed with a MINFLUX acquisition of VGlut (localizations in transparent violet). VGlut is labeled by AF647 with primary and secondary antibody. Clusters assigned by analysis with the dbscan algorithm are fitted with a circle and the circle is overlayed. **C** Results of the cluster analysis of VGlut. **D** AMPA receptor structure from PDB file 3KG2 and extracted distances between the labeled amino acids. **E** MINFLUX acquisition of AMPA receptors directly chemically labeled with AF647. Localizations from the same emission event and localizations that fall within 2 nm of each other are assigned to the same molecule (AMPAR subunit). Molecules are plotted as cyan dots. **F** Zoom-in to the image region with highest molecule density (putatively the post synapse) and cluster analysis of the AMPA receptors. Clusters are found by the dbscan algorithm and their border is delineated using a spline-fit. **G** Cluster analysis results of AMPA receptors. **H** Distances between AMPAR subunit localizations (nearest neighbors, second nearest neighbors and third nearest neighbors). Loc. – localizations. d_NN_ – distance to nearest neighbor(s).

Piccolo, a protein situated close to the synaptic release site, served as context for a MINFLUX acquisition of VGlut (Fig. 5B). In these experiments, VGlut was labeled by Alexa Fluor 647 with primary and secondary antibody. The apparent clustering was quantified further, with clusters assigned by analysis with the dbscan algorithm and shown by overlayed circles. Results of the VGlut cluster analysis (Fig. 5C) indicated a median diameter of 39 nm and containing ∼28 localizations on average.

The AMPA receptor distribution was visualized by direct labeling with a chemical AMPAR modification (CAM2) probe conjugated to Alexa Fluor 647 (33). The probe labels three out of four possible AMPAR subunits, specifically GluA2, GluA3 and GluA4, but not GluA1. Functional AMPA receptors are tetramers, composed differently from GluA1-GluA4. Homogenous AMPA receptors containing only one subunit type and mixed AMPARs containing several subunit types are possible (34, 35). Up to a maximum of 4 sites (subunits) could therefore be labeled. From structural data of the AMPA subtype ionotropic glutamate receptor (PDB 3KG2,(36)), the spatial arrangement of the amino acid in the AMPAR targeted by labeling was extracted for reference (Fig. 5D).

In analyzing the MINFLUX recordings, localizations from the same emission event and localizations that fall within 2 nm of each other were assigned to the same AMPAR subunit (Fig. 5E). The image region with highest molecule density, putatively the post synapse, was subjected to a cluster analysis by the dbscan algorithm, and cluster borders were delineated using a spline-fit (Fig. 5F). AMPAR clusters featured a median cluster area of ∼60 nm^2^ and approximately 7 (median) fluorophores (i.e. labeled AMPAR subunits) present per cluster (Fig. 5G). Intriguingly, the distances between AMPAR subunit localizations – the average nearest neighbors, second nearest neighbors and third nearest neighbors (histogram peaks at ∼4 nm, ∼5 nm, ∼7 nm) quantified across the entirety of subunit positions in the field of view – were found to correspond approximately with the distances extracted from structural data (Fig. 5H). A comparison with simulation of AMPAR subunit localization yielded similar nearest neighbor distributions (compare SI Materials and Methods, and Fig. S9). This shows that, while the sampling of individual localization patterns may have been incomplete in many cases, the ultra-precise nature of MINFLUX localization rendered statistically valid mutual distances even in tissue. In fact, these distances were quite consistent with the intra-molecular spacings of expectation. For instances of collections of sufficient points, individual patterns illustrate how trapezoidal arrangements may be annotated (Fig. S10).

### 3D MINFLUX scheme for tissue imaging

As only a subset of dendrites was labeled, we expected to observe PSD95 clusters whose dendrites were not labeled. With ∼600 nm extent projected along the optical axis in the 2D MINFLUX imaging without depth discrimination, there was a clear potential for misclassification of PSD95 proteins that appear to localize on the dendrite but could also be positioned above or below. Quantification of distances among proteins in 2D is inherently prone to projection errors. We therefore chose to explore 3D localization for further two-color studies.

In prior reports of the first 3D MINFLUX demonstrations (14, 15, 17), a ‘z-doughnut’ which forms an intensity minimum in three dimensions was used for localization. The intensity gradient of such a ‘z-doughnut’ is inevitably stronger along the optical axis than in the perpendicular direction. Further, the minimum (zero) definition contrast, especially perpendicular to the optical axis, decreases rapidly if there are sample-induced aberrations (9) (compare Fig. S4, Supplementary Information). The zero definition is laterally considerably less robust (deep) in tissue than the maximum of the regularly focused beam and the zero of the doughnut (Fig. 6A). Transverse intensity profiles through simulated focal distributions illustrate this effect with increasing spherical aberration strength.

**Fig. 6.**
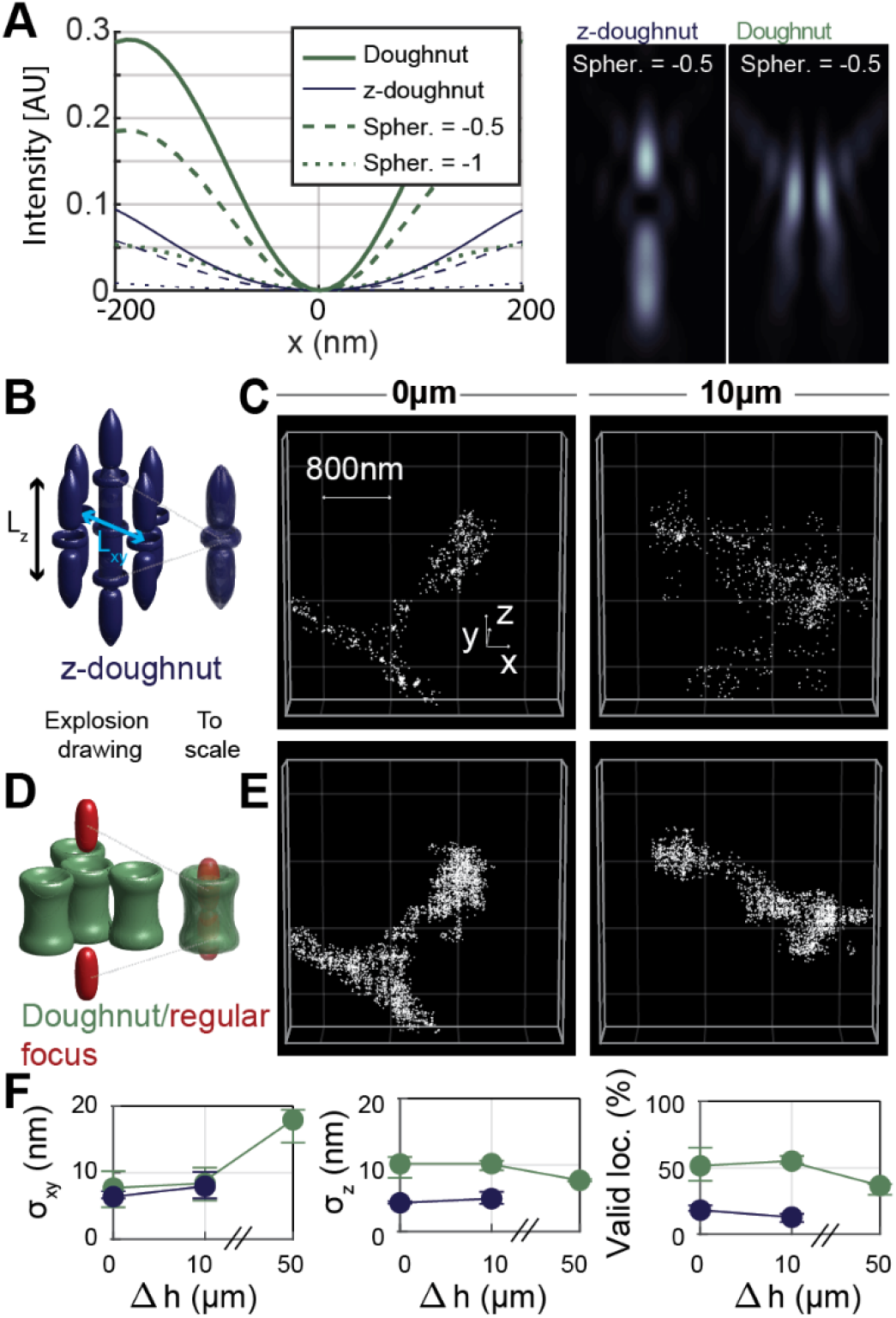
3D MINFLUX localization scheme for imaging in aberrant media. **A** Simulated focal intensity profiles for different spherical aberrations. While the z-doughnut provides good imaging contrast along the optical axis, its contrast perpendicular to the optical axis was notably worse than for the doughnut, especially in the case of spherical aberrations. **B** Standard 3D MINFLUX localization scheme based on the z-doughnut. **C** 3D MINFLUX images of spines in 0 and 10 µm imaging depth in tissue acquired with the z-doughnut. **D** Doughnut/regularly focused beam 3D imaging scheme. **E** Same regions of interest as in B acquired with the Doughnut/regular focus imaging scheme. (Interleaved acquisition: first z-doughnut, then doughnut/regular focus, then again z-doughnut). **F** MINFLUX metrics for 3D imaging with doughnut/regular focus or z-doughnut in different imaging depths. Spher. – first order spherical aberration (Zernike coefficient), AU – arbitrary units, loc. – localizations, σ – localization precision.

In the absence of tissue homogenization or clearing methods, we chose to implement and explore a new 3D localization pattern consisting of two excitation beam shapes: the regular focus for localizing along the optical axis and the doughnut (‘vortex’) beam for localizing perpendicular to it. This 3D localization pattern was utilized for the last iteration step of the iterative MINFLUX scheme. The previously described z-doughnut-based iterative MINFLUX scheme and our new doughnut/regular focus-based iterative MINFLUX scheme are illustrated in Fig. 6B and Fig. 6D.

Choices of parameters of the modified least mean squares (mLMS) estimator for the z-doughnut localization were guided by previous work (14) and adapted for a bigger localization range along z (detailed in Table S5 (*Supplementary Information*)). For the doughnut-regular focus scanning pattern, mLMS estimator parameters were found by repeatedly localizing the center of mass of fluorescent beads with the calibrated setup, displacing them by known distances with the fine stage and checking that the estimated center of mass of the fluorescent beads moved by the expected distance.

We compared the performance of our new 3D MINFLUX scheme to the ‘z-doughnut’ by interleaved acquisition, observing higher valid-event rates on the same structure in dendritic spine actin samples (Fig. 6E vs. 6C). The lateral precision was comparable, whereas axial precision was about half (∼10 nm) of the precision achieved with the *z*-doughnut. Valid localization rates were improved several-fold, enabling robust imaging (Fig. 6F). The results experimentally confirm the reasoning that while the z-doughnut provides rather good imaging contrast along the optical axis, in the presence of aberrations, the contrast in the lateral (*xy*) dimensions becomes notably worse than for the lateral doughnut.

### Investigation of PSD95 and AMPA receptor distributions in 3D on the post synapse

3D two-color MINFLUX (Fig. 7) enabled the mapping of protein localizations as 3D spatial distributions in context: we illustrate this with PSD95 along with actin on the post-synapse (Fig. 7A,D), or with the distribution of AMPA receptors together with the distribution of PSD95 (Fig. 7C).

**Fig. 7.**
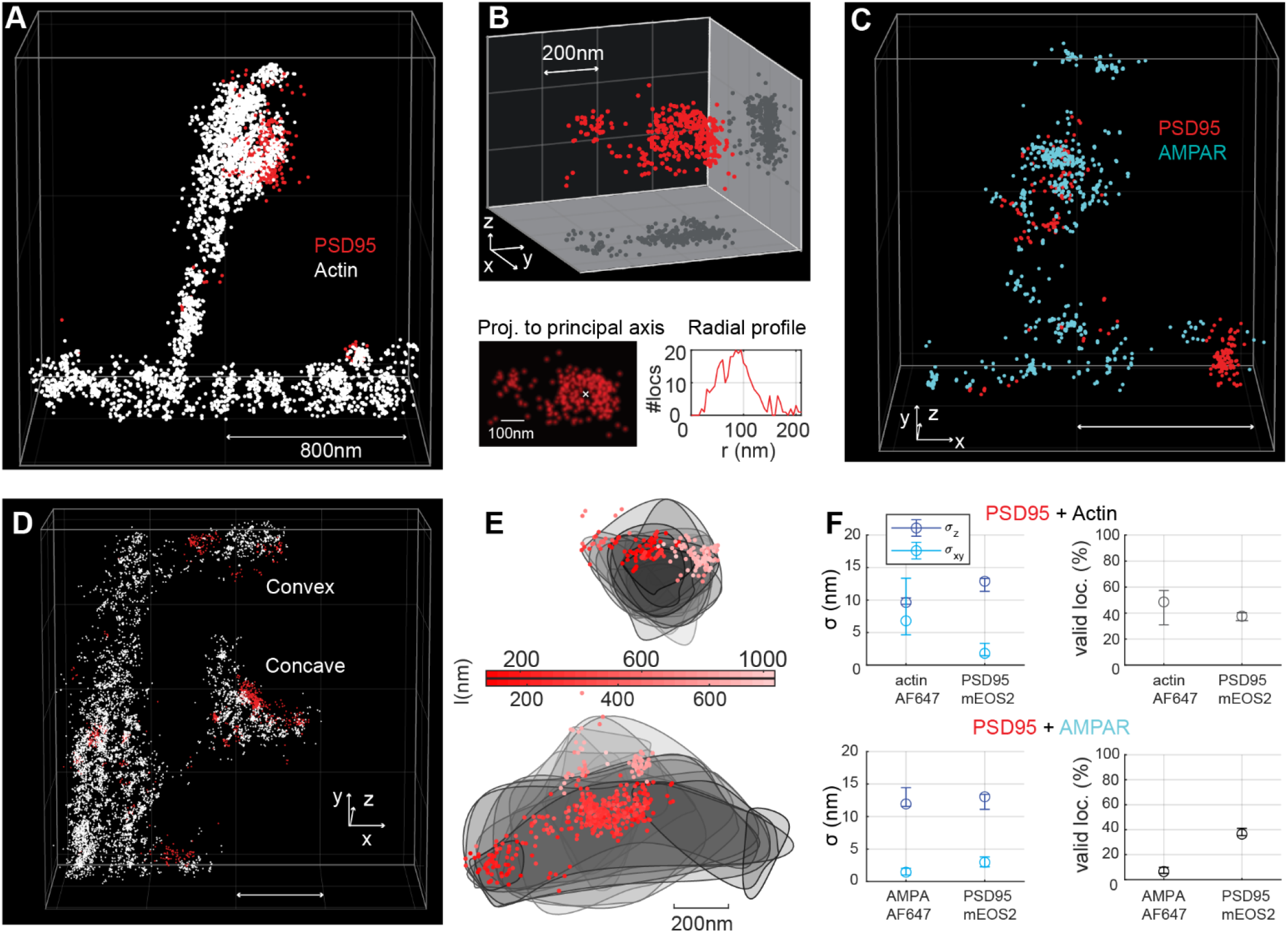
3D MINFLUX with two excitation colors for investigating the protein distributions on the post synapse (spine head). **A** Region of interest showing part of a dendrite with a spine. Spine morphology is delineated similarly to the 2D two-color MINFLUX recording of PSD95/actin for all images shown (**A, C** and **D**). Localizations plotted as scatter with marker size of 15 nm (∼3σ). Localizations of actin are plotted in white and of PSD95 in red. **B** PSD95 distribution on the spine head shown in **A**. The PSD95 proteins are located mainly on one side of the convex spine head. Notably, the PSD95 proteins are clustered and the biggest cluster is perforated. Projection of the PSD95 localizations to an orthonormal head-on view, and radial profile centered on the hole in the PSD95 cluster. **C** PSD95 and AMPA receptor distribution on a spine with similar morphology as in **A**. AMPA receptor localizations are plotted in blue. The highest AMPA receptor density is on the top of the convex spine head. **D** Region of interest containing two spines: one with a concave-shaped spine head and one with a convex-shaped spine head. **E** Convex and concave spine head in **D**, projected from the top of the spine head. Gray lines delineate 80 nm thick slices of actin and PSD95 localizations. PSD95 localizations shown as scatter, with the distance l along the line of sight from the top of the spine color-coded. **F** MINFLUX metrics over all 3D two excitation color MINFLUX images of actin and PSD95 as well as AMPAR and PSD95. Shown are median values, whiskers represent the interquartile range. l – length along the viewing axis, Proj. – projection, loc. – localizations, σ – localization precision. Animations of the data in A,C and D are found in Movies S1-S3. Scale bar: 800nm (**A, C, D**) and as indicated.

PSD95 arrangements may be analyzed in terms of their spatial extent and internal distribution. This is exemplified by a large group of PSD95 that sits in a rather large (∼200 nm) diameter cluster which exhibits a central perforation (void) (Fig. 7B), as suitable projection of the 3D dataset and radial distribution analysis (maximal radius ∼90 nm) shows.

The complex distribution of the directly-labeled AMPA receptors was visualized as a further illustration of 3D capabilities in conjunction with PSD95. The AMPAR protein number density was appreciably highest at the top of the convex-shaped spine head (indicated by white arrow).

Spine heads of different morphology, such as convex and concave shapes, were appreciable in imaged examples based on the volumes outlined by actin (Fig. 7D). PSD95 accumulations can be related in space to the underlying spine head geometry. For instance, a prominent PSD95 cluster sat distinctly on the side of a convex, bulb-shaped spine head (Fig. 7D,E upper example), whereas the clustered PSD95 sat centrally on an elongated concave spine head (lower example in Fig. 7D, with oriented display of data in Fig. 7E).

With respect to quantitative performance, spatial precisions fell in the range of ∼10-12 nm axially (Fig. 7F), and well below (∼2-7 nm) in the lateral directions. The direct attachment of Alexa Fluor 647 to AMPAR led to shorter emission bursts and less robust localization success rates (lower valid localization fraction). At the same time, repeated on-blinks from individual copies of the dye likely contributed to a putatively rather complete sampling of the present dye molecules over the whole recording duration, and the events that were localized exhibited the usual high MINFLUX precision (Fig. 7F).

## Discussion and Conclusion

It goes without saying that any fluorescence microscopy, including MINFLUX, records nothing but the fluorophores. This obvious fact becomes highly relevant at the own scale of the proteins. Therefore, any conclusion drawn from the detection and localization of fluorophores has to take this fundamental limit of fluorescence imaging into account. Nonetheless, our initial 2D and 3D, two-color MINFLUX results in tissue, with <10 nm 3D fluorophore localization, open up entirely new avenues to investigate protein distributions on the single-synapse level in fixed and living brain slices. Synaptic heterogeneity at the molecular level in intact tissue contexts (28, 37) may be assessed with resolution at the few-nanometer scale, especially with forthcoming developments of (semi-)automated acquisition strategies that allow to select or automatically identify ROIs based on experiment criteria.

The fact that single-digit nanometer localization precision is reached in the complex tissue environment is arguably remarkable. In this context, it is worth commenting that close-by fluorophores in the sample involve virtually identical optical paths through the sample and hence basically identical optical conditions, such as sample inhomogeneities. Any potential aberration-related spatial shifts in the minimum of the MINFLUX excitation pattern (which would lead to a shift in the spatial position of the recorded fluorophore) therefore affect close-by fluorophores identically. Thus, a faithful mapping of the relative position of adjacent fluorophores in the optically inhomogenous sample is ensured. This is why in MINFLUX nanoscopy the resulting nanoscale distributions of the resolved molecular localizations can be interpreted with high confidence.

Our demonstrations further indicate that the use of a silicon oil immersion objective lens enables this MINFLUX resolution without dedicated online aberration corrections to respond to sample-dependent depth aberrations. The absence of suitable bright reference objects (‘guide stars’) at the required depth – which would be needed to perform an optimization of the excitation pattern at each new ROI – would make this more challenging to realize. Instead, the described results were obtained by setting up the optics based on calibration measurements at the coverslip surface, with no further adjustments at depth.

Nonetheless, looking ahead, the prospects for the development of online aberration correction highlight a further attractive feature of MINFLUX in this regard. In PALM/STORM, the localization relies completely on a large but inherently limited number of emitted fluorescence photons. This full reliance on emitted fluorescence makes it challenging to implement effective aberration correction methodology, as any aberration correction approach inherently uses up emitted photons as well. In contrast, aberration correction in MINFLUX can be carried out by modifying the shape and direction of the stable and bright laser beam rather than the feeble fluorescence ‘beam’ delivered by the fluorophore. The alternative but unperfect remedy of using an auxiliary laser beam for aberration correction would also not be needed.

As a point of particular interest, and as our PSD95 imaging demonstrates, MINFLUX imaging in tissue is also realized with fluorescent proteins, for which the number of emitted photons is more limited than for organic dyes but which are better compatible with living tissue. FP fusion proteins also inherently maintain a stoichiometric mapping of protein copy numbers (1:1 correspondence) in the microscopy analysis. An additional advantage of MINFLUX imaging in biological tissue is the need for comparatively low laser powers and only local illumination, which should be more compatible with imaging of living tissues over long periods of time.

## Author Contributions

S.W.H. recognized the potential of MINFLUX to image deep in scattering samples and outlined the research project with T.M., V.W. and J.K.P., who designed and built the MINFLUX setup for tissue imaging with input by S.W.H. T.M. and V.W. evaluated the setup performance. T.M. and K.A.K. designed and iteratively improved the experiments. T.M. performed the MINFLUX measurements. K.A.K. suggested the samples to be investigated, provided the tissue samples and carried out the tissue stainings. K.A.K. and H.S. prepared the tissue samples. M.L. calibrated the deformable mirror, developed electronic hardware and FPGA software for its real-time control, and advised on PSF calculations. S.G.N.G. provided the transgenic PSD95-mEOS2 mouse line. T.M., K.A.K. and V.W. analyzed the data. T.M., K.A.K., S.J.S. and S.W.H. wrote the manuscript. S.J.S. and S.W.H. gave critical feedback on the results during the project.

## Acknowledgement

We thank Ellen Rothermel for cell stainings and Tanja Koenen for providing and maintaining cell lines. Michael Weber for helpful discussions about caveolae. Jan Seikowski (Facility for Synthetic Chemistry, MPI-NAT) synthesized the AMPAR probe, in coordination with Vladimir Belov (NanoBiophotonics, MPI-NAT). Jochen Staiger and Patricia Sprysch supported handling of the vibratome for fixed slices preparation. Theocharis Alvanos and Shalini Pradhan helped with the living slices preparation (living slices vibratome). We also thank the team at the animal house for support with the mouse lines, Marco Roose for IT support and a custom solution for SLM control, the precision mechanics workshop (MPI-NAT), especially Mario Lengauer, and Rainer Pick for manufacturing custom components and the precision optics workshop, especially Matthias Kulp (NanoBiophotonics, MPI-NAT), for custom optics solutions. T.M. is part of the Max Planck School of Photonics supported by BMBF, Max Planck Society, and Fraunhofer Society. S.G.N.G. was supported by the European Research Council (ERC) under the European Union’s Horizon 2020 research and innovation programme (695568 SYNNOVATE). Fig. 1 B, C was created with BioRender.com.

