## Supplementary Information for "MINFLUX fluorescence nanoscopy in biological tissue"

**Contents:**

Materials and Methods
Supporting text
SI References
Figures S1 to S11
Tables S1 to S10
Legends for Movies S1 to S3

**Other supporting materials include:**

Movies S1 to S3

#### Materials and Methods

##### Animals

Animal procedures described here were carried out in accordance with institutional regulations on animal use in research. Experiments performed on living animals were approved and authorized by the Lower Saxony State Office for Consumer Protection and Food Safety (Niedersächsisches Landesamt für Verbraucherschutz und Lebensmittelsicherheit (LAVES)). Sacrificing rodents for subsequent preparation of living slices and cultures did not require specific authorization or

notification (Animal Welfare Law of the Federal Republic of Germany Tierschutzgesetz der Bundesrepublik Deutschland (TierSchG)).
All mice were housed with a 12 hours light/dark cycle and an *ad libitum* access to food and water.

#### Stereotaxic Injections

The protocol of stereotaxic injection has been described previously in (1, 2). Briefly, the adult mouse of C57BL/6J background was anesthetized by 1.0-2.0 % isoflurane (Isofluran CP, CP Pharma) in oxygen-enriched air (47.5 % oxygen, 50 % nitrogen, and 2.5 % carbon dioxide; Air Liquide) and fixed into a stereotaxic frame (SG-4N, Narishige). The scalp was incised and a craniotomy was performed on the parietal bone above the visual cortex of the left hemisphere. A pre-pulled, tapered, borosilicate glass injection capillary (World Precision Instruments, cat. 1B150F-4) was filled with a solution of pAAV-hSyn-Lifeact-EYFP virus (3) diluted 1:5 in sterile artificial cerebrospinal fluid (ACSF; NaCl 126 mM, KCl 2.5 mM, $CaCl_2$ 2.5 mM, $MgCl_2$ 1.3 mM, HEPES 27 mM, glucose 30 mM; pH 7.4). The capillary was subsequently lowered ca. 500 µm into the brain with an angle of 20° to the horizontal axis. A volume of 250–500 nl of virus-containing solution was injected using a pressure application system (TooheySpritzer, Toohey Company) generating 30 ms pulses delivered with 20 psi on manual command. The capillary was retracted and the scalp was surgically stitched by polyamide surgical suture (6.0, Ethilon, cat. 697H). Subsequently, the animal was allowed to recover from anesthesia.
Perioperative analgesia was achieved by subcutaneous (s. c.) injection of Buprenorphine (Buprenovet, Bayer) and local analgesia of incision sites by s. c. injection of 2 % Lidocaine (Lidocainhydrochlorid 2 %, Bela-Pharm). Throughout the surgery, eyes were protected from dehydration by application of ointment (Bepanthen, Bayer) and a custom-built heating plate was used to maintain mouse body temperature. Analgesic and anti-inflammatory post-surgical care was achieved by s. c. administration of Carprofen (Rimadyl, Zoetis).

#### Intracardial perfusion fixation and fixed brain slice preparation

Intracardial perfusion took place following the period of around 4 weeks from the time of stereotaxic injection. Injected mice were anaesthetized by intraperitoneal injection of an overdose of Ketamine (Ketamin 10 %, Bela-Pharm) and subsequently transcardially perfused with Phosphate Buffered Saline (PBS, pH 7.4), followed by 4 % PFA in PBS (pH 7.4). The brain was dissected and post-fixed by an overnight incubation in 4 % PFA/PBS at 4°C. Afterwards, the fixative was removed, the brain was transferred into PBS (pH 7.4) and sliced into 60-120 µm thick consecutive coronal or sagittal sections using a vibratome (VT1200S, Leica).

#### Immunohistochemistry

Fixed slices were successively washed two times with Tris Buffer (TB), Tris Buffer Saline (TBS) and Tris Buffer Saline containing 0.5 % Triton-X 100 (TBST) (all with pH 7.6) for 15 min each at room temperature (RT). The slices were subsequently blocked with 10 % normal goat serum (NGS; Jackson ImmunoResearch, cat. 005-000-121) and 0.25 % bovine serum albumin (BSA) in TBST (blocking solution) for 1.5 h at RT. The tissues were then incubated with primary antibodies diluted in blocking solution for 72 h at 4$^{o}$C on a rocking plate (primary antibodies used here: either anti-GFP (dilution 1:300; Abcam, cat. ab6556), anti-Caveolin-1 (dilution 1:100, Cell Signaling, cat. 3267) or anti-VGlut1 (dilution 1:200; Synaptic Systems, cat. 135 308). Additionally, slices labelled with anti-VGlut1 antibody were co-stained with anti-Piccolo antibody (dilution 1:300, Synaptic Systems, cat. 142 104). The slices were washed 4 times with TBST for 15 min each at RT and stained with Alexa Fluor 647-conjugated goat anti-rabbit secondary antibody (Thermo Fisher Scientific, cat. A-21245) diluted 1:1000-1:10000 in TBST for 4 h at RT on a rocking plate. Slices co-stained with anti-Piccolo antibody were additionally labelled with Alexa 532-conjugated donkey anti-guinea pig secondary antibody (Dianova, cat. 706-005-148) diluted 1:500 in TBST. Slices were successively rinsed 2 times with TBST, TBS and TB, and stored at 4$^{o}$C in TB buffer until nanoscopy.

#### Injection of beads into fixed brain slices

A volume of 250-500 nl of PBS solution containing TetraSpeck fluorescent beads with average size of 100 nm (dilution 1:200; Thermo Fisher Scientific, cat. T7279) was injected into the 120 µm-thick, fixed coronal brain slice using ToohеySpritzer pressure application system. The bead solution was injected into several locations to cover the full depth of the slice.

#### Preparation of agarose-sucrose-bead sample

Agarose-sucrose pads were prepared similarly to the method described in (4). In short, a volume of 750 µl of distilled water was mixed with 8-24 mg agarose (low gelling temperature; Sigma-Aldrich, cat. A6560-25G) and 318-680 mg sucrose. The whole sample was weighted. Under constant stirring, the sample was warmed up to 100°C, held at this temperature for 2 min, then cooled down and held at 80°C for 2 min. Distilled water was added to fill up to the previous weight of the sample. Then, a volume of 250 µl of 100 nm size TetraSpeck fluorescent beads (Thermo Fisher Scientific, cat. T7279) diluted 1:200 in water was added to the sample while stirring on the heating plate. A cavity slide and a coverslip (thickness: 170±5 µm) were heated up to 80°C. A volume of around 70 µl of prepared solution was pipetted into the cavity simultaneously sliding the coverslip over to avoid air bubbles in the sample. The rest of the solution was poured onto a slide, cooled down to RT, detached from the slide and used for refractive index measurement with the refractometer (Schmidt + Haensch, ATR-L).

#### Preparation of living brain slices

Living slices were prepared of brains isolated from adult PSD95-mEos2 C57BL/6J mice of both genders (5). Mice were sedated with isoflurane in a sealed container and quickly euthanized by cervical dislocation. Brains were dissected instantly, transferred into ice cold ACSF (pH 7.4) infused with 95 % $O_2$ and 5 % $CO_2$ and sliced into 300 µm thick coronal sections using a vibratome straight away. Prepared slices were either stained with CAM2-Alexa Fluor 647 compound (6) or directly imaged with MINFLUX nanoscope.

#### Chemical labeling of AMPA receptors

The protocol of chemical labeling of AMPA receptors was adapted from (6). In brief, living brain slices were stained with 1 µM CAM2-Alexa Fluor 647 in ACSF (pH 7.4) infused with 95 % $O_2$ and 5 % $CO_2$ for 1h at RT on a rocking plate. The slices were then washed 3 times with ACSF and fixed for 3 h with 4 % PFA in PBS (pH 7.4) immediately after washing. Fixed slices were rinsed 3 times with PBS (pH 7.4) and stored at 4°C until nanoscopy.

#### Synthesis of AMPAR modification (CAM2- Alexa Fluor 647 conjugate)

CAM2-Alexa Fluor 647 conjugate was synthesized following the method described in (6) with minor modifications.

#### Cells

U-2 OS cells (ECACC, cat. 92022711, lot 17E015) were cultured in modified McCoy's 5A medium (Thermo Fisher Scientific, cat. 16600082) supplemented with 10 % (v/v) FBS (Bio&Sell, cat. S0615), 1 % (v/v) Sodium Pyruvate (Sigma, cat. S8636) and 1 % (v/v) Penicillin-Streptomycin (Sigma, cat. P0781) in a humidified 5 % $CO_2$ incubator at 37°C. Cells were seeded on coverslips 24 h before staining.

#### Immunocytochemistry

U-2 OS cells were washed with PBS and subsequently fixed with 8 % (w/v) PFA in PBS for 5 min at 37°C. Cells were then permeabilized with 0.5 % (v/v) Triton-X in PBS for 5 min at RT and blocked with 2 % (w/v) BSA in PBS (blocking solution) for 10 min at RT. Afterwards, cells were incubated with primary rabbit anti-Caveolin-1 antibody (Cell Signaling, cat. 3267) diluted 1:200 in blocking solution for 1h at RT. Subsequently, cells were rinsed with blocking solution and incubated with Alexa Fluor 647-conjugated goat anti-rabbit secondary antibody (Thermo Fisher Scientific, cat. A-21245) diluted 1:5000 and 1:7000 in blocking solution for 1h at RT. Cells were then washed with PBS (pH 7.4) and stored at 4ºC until nanoscopy.

#### MINFLUX nanoscope for tissue samples

A MINFLUX nanoscope for imaging in tissue was built, based on the nanoscope design described in (7, 8). A detailed overview of the confocal beam-scanning MINFLUX setup, is given in Fig. S2. The nanoscope is equipped with a silicon oil objective (*NA*=1.35; UPLSAPO100XS, Olympus) and a fully computer-controlled sample positioning system. The sample positioning system consists of a coarse (50nm step size) stepper-motor z-stage (M-230.25, Physik Instrumente (PI)) with a LVDT inductive sensor for measuring displacement along the optical axis, a coarse (100 nm positioning resolution) xy-stage (M-686.D64, PI) and a fine (0.1 nm positioning resolution) piezo-driven xyz-stage *(*P-733.3DD, PI). An imaging-depth adaptable lock system (described in detail in main text, Fig. 2) for stabilizing the 3D sample position on the nanoscope was built in close proximity to the objective.

Excitation and activation lasers with their ventilation and their power modulation are mechanically decoupled from the main setup (see Fig. S2D). The setup contains three excitation lasers (2RU-VFL-P-2000-647-B1R and 2RU-VFL-P-5000-560-B1R, MPB Communications; IBEAM-SMART-511-S-HP**,** Toptica) and the activation laser (Phoxx 405-60, Omicron) with corresponding power modulation via acousto-optical modulators (AOMs, (MT110-A1-VIS)) and acousto-optic tunable filters (AOTFs, (AOTFnc-VIS-TN, AA Opto-Electronic)) and spectral selection possibility (AOTFs). The Toptica laser with integrated power modulation does not require an AOM. The three excitation laser beams are rearranged and coupled into the PM fibers for the red (647nm) and the green (511 or 561 nm) doughnut excitation beam path, the regular focus excitation beam path (all three wavelengths possible) and the widefield excitation beam path (all three wavelengths possible). Activation light can be either added to the widefield beam path or separately via the activation beam in-coupling, allowing for a regular focus activation beam in the sample plane.

The excitation beams, starting from the polarization maintaining fiber (PM) -outcoupling (separate paths for the regularly focused beam and the (z-) doughnut beam), can be phase modulated by the spatial light modulator (SLM, (SLM-100*,* Santec)). The phase modulation by the SLM allows beam shaping into the (z-)doughnut in two separate spectral channels and aberration correction. From the SLM, the laser light passes the beam scanning system and enters the objective. The beam scanning system consists of electro-optical deflectors (EODs, (M-311A AD*P, Conoptics)) for fast beam scanning over a small area perpendicular to the optical axis (xy), the Tip/tilt piezo mirror (TM, (PSH 10/2 SG, piezosystem jena)) for (de-)scanning a bigger field of view (in xy) and the deformable membrane mirror (DMM, (Multi-3.5, Boston Micromachines Corporation)) for fast z-(de-)scanning. Several 4f-relay lens systems adapt the beam diameter and relay the scanning system pupils into the back aperture of the objective lens. Activation light is coupled-in after the EODs. A dichroic mirror (DM, (zt440/514/561/640rpc, Chroma)) reflects the activation and excitation wavelengths towards the objective and transmits the fluorescence emission to the detection. A motorized DM (ZET405/514/647 TIRF, AHF) placed between DMM and objective, allows the switching from the confocal or MINFLUX imaging mode to the widefield imaging mode by inserting the DM into the main beam path. Motorized achromatic quarter- and halfwave- plates (mQWP, mHWP, (RAC 3.2.15, RAC 4.4.15, B. Halle)) were used to control the polarization of the excitation light in the back aperture of the objective.

The first detection box, directly behind the main DM, contains notch filters for blocking light from the excitation and activation lasers (ZET647NF, ZET561NF, ZET405NF Chroma and ZET514TopNotch, AHF) and a short pass filter (ET750sp-2p, Chroma) for blocking light from the focus lock system. Motorized flip-mirrors are used to direct the fluorescence either onto the widefield camera, the large area APD (LA APD) for PSF measurements or the confocal pinhole with variable size for spatial filtering and then onto the multimode fiber leading towards the multicolor detection. The large area APD has a detection area of >> 1 AU (non-confocal APD).

The fluorescence light that passes through the first detection box (subpanel Fig. S2 C) and is selected by the variable confocal pinhole (MPH16, Thorlabs) is coupled into a multimode fiber (M50L02S-A, Thorlabs) and coupled-out in the Multicolor-Detection (Subpanel Fig. S2 B). In the Multicolor-Detection, fluorescence light is spectrally separated into a green, an orange and a red spectral channel by DMs (FF560-FDi01, FF640-FDi01, Semrock). Fluorescence light in the three resulting spectral channels, is split by customized tunable filters (TSP-01-561-25x36, TSP-01-625-25x36, TSP01-704-25x36, Semrock), that can be spectrally tuned by computer-controlled servo motors, to a custom-chosen ratio and imaged onto the detection avalanche photodiodes (APDs, (SPCM-AQRH-13-TR, Excelitas Technologies)). For good isolation against ambient light, two layers of light-proofing boxes are built around the multicolor-detection box. The rest of the setup is housed in with one light-proofing layer.

#### Nanoscope control software

We adapted the custom written LabView programs from previous MINFLUX implementations (7, 8). The focus lock software was augmented to include the control of the adaptive lens and the absolute positionable mirror. A control system for the servo motors moving the half-wave plate and the quarter-wave plate was added to the main MINFLUX user interface. This allows to optimize the circular polarization for different excitation colors. A new 3D localization pattern was implemented, combining regular focus and doughnut exposures. A functionality to ramp-up the activation intensity was added (described in detail in the main text, Fig. 2).

A control unit for the DMM for focus-scanning consisting of an FPGA (PCIe-7852R, National Instruments) and a Camera-Link interface was linked to the main FPGA (USB-756R, National Instruments). The host program for user interface to the DMM FPGA and the code for the DMM FPGA were written in LabView (LabView 2020 sp1, 32bit).

#### MINFLUX data acquisition

Data were acquired following the steps described in (8) with some adaptations. The widefield fluorescence image was used to choose the brain region of interest (ROI). Due to the better optical sectioning in confocal mode, the ROI for MINFLUX acquisition was selected based on confocal overview scans. Because Alexa Fluor 647 is in an emitting state before starting the MINFLUX measurement, the structure of interest can be seen by acquiring a confocal or widefield image before the MINFLUX measurement. In the case of the fluorescent protein mEOS2, that photo-converts from the bluer to a redder emitting state upon illumination with 405 nm light, the ROI was selected by imaging the blue emitting form.  Before MINFLUX acquisition, Alexa Fluor 647 was switched-off by applying excitation light at 647 nm. Activation and probing of the fluorophore emission were started by scanning with steps of 150-200 nm to activate or photo-convert fluorophores within the ROI homogenously. Emitting molecules were iteratively centered and localized with the modified least mean squares (mLMS) estimator (as described in (7)) implemented on the FPGA. The variable pinhole was used to adapt the confocal pinhole size to optimize the MINFLUX metrics in the sample. For the statistics shown in Fig.4 C-E, statistics are pulled together from the pinhole range 0.4-0.6 Airy Units (AU).

Sample-dependent measurement parameters for 2D MINFLUX are detailed in Table S4. For 3D MINFLUX acquisitions, parameters are shown in Table S6 and S7. In the z-doughnut vs. doughnut/regular focus acquisition scheme comparison, the MINFLUX image employing the z-doughnut was acquired first, then the doughnut/regular focus scheme was run over the same field of view, then again, the z-doughnut scheme was used. Data sets with z-doughnut were combined.

#### Progressive activation

The imaging time per molecule can be divided into three components: (1) $t_{act}$, the duration to activate the molecule, (2) $t_{readout}$, the duration to read-out the photons from the activated molecule and (3) $t_{jump}$, the time to move to the next location. (1) and (2) are repeated until either no further activations are detected for a maximal duration $t_{act,max}$ or a maximum imaging time $t_{image,max}$ has been reached at the current location.

Typical values are: $t_{act,max}$ = 2 s, $t_{readout}$ = 0.1 – 1 s, $t_{jump}$ = 200 ms.

$t_{readout}$ cannot be further reduced due to the maximal photon flux that can be extracted from the molecule. Meanwhile, $t_{jump}$ is due to the response time of the tip-tilt piezo mirror in the system. The probability to activate depends linearly on the dose of activation light applied and the density of molecules that can be activated within the activation volume. The activation intensity has to be low enough that even in regions of high densities of activatable molecules, the probability that more than one molecule is activated at the same time is low. This leads to a typical activation scheme where the activation intensity starts out low and is increased manually when the activation rate appears to be too low after the whole field of view has been imaged repeatedly and the density of activatable molecules had been reduced significantly.

This approach has drawbacks:

1. in regions with a low molecule density, $t_{act}$ is very long and can often reach $t_{act,max}$
2. the user has to intervene constantly by increasing the activation power to keep the average activation rate reasonably high.

Therefore, we implemented a new activation scheme that we are calling Progressive Activation: It starts with a brief period (e.g. 20 ms) of almost no activation (limited by contrast of the digital laser power modulation), to avoid additional activations in case a molecule is already activated. Then it continues at a low activation intensity (about 2 % of the final intensity) that ramps up exponentially to the maximum intensity, see Fig. S11 A. The reason for the exponential increase is that for each time interval the activation probability is increased by the same factor. The time needed for constant (or homogenous) activation to reach the same activation dose as progressive activation is estimated. Within 0.3 s the Progressive Activation would have applied the same activation dose as the homogeneous activation within more than 2 s, see Fig. S11 B,C.

These means in turn that the $t_{act,max}$ of 2 s of activation previously needed at a given spot can be compressed into 0.3 s; and, if fluorophores to be activated are present, to an even shorter period of time.

#### Sample mounting

To stabilize the sample during the MINFLUX measurement, all coverslips used for mounting were pre-treated with gold nanorods (Nanopartz, cat. A12-25-850-CTAB-DIH-1-25) employed here as fiducial markers. The nanorods were applied as described in (8). In brief, nanorods were diluted 1:3 in PBS, sonicated for 10-15 min at RT and incubated onto a high precision coverslip (170±5 µm thick) for 10 min at RT. Afterwards the coverslip was rinsed several times with PBS to remove unbound nanorods.

Anaerobic redox blinking buffer containing 0.8 mg/ml glucose oxidase (Sigma-Aldrich, cat. G2133), 128 µg/ml catalase (Sigma-Aldrich, cat. C100-50MG), 50 mM Tris-HCl (pH 8), 10 mM NaCl, 10% (w/v) glucose and 30-50 mM MEA (cysteamine hydrochloride; Sigma-Aldrich, cat. M6500) was used for MINFLUX imaging of fixed samples stained with Alexa Fluor 647, following the procedure described in (8). Two-color images of Alexa Fluor 647 and mEOS2 were also acquired using this buffer.

Fixed brain slices with PSD95 endogenously fused with the photoconvertible fluorescent protein mEOS2 were imaged in 50 mM Tris buffer (pH 8) in 95% $D_2O$ to reduce the short time blinking and increase the photon count of mEOS2 (9).

All fixed samples were sealed with dental glue (eco-sil speed, Picodent, cat. 1300 7100).

Living brain slices were imaged in a custom-built imaging chamber shown in Fig. S1. Briefly, 300 µm thick living slices were placed onto a high precision coverslip pre-coated with nanorods and pre-fixed to the chamber with eco-sil speed dental glue. Slices were secured by a plastic grid magnetically attached to the imaging chamber and imaged in ACSF infused with 95% $O_2$ and 5% $CO_2$.

#### MINFLUX data analysis for depth imaging series

Trace segmentation and position estimation were performed as described in (8), with the adaptation, that the starting threshold for estimating signal emission was set automatically in the depth imaging measurement series. For automatic estimation of the starting threshold for signal emission, a quantile filter was set on the emission trace in the last iteration that removes extreme outliers. Then, the remaining emission trace was fitted to a mixture of two Gaussians and the threshold between them was taken as optimal threshold between signal and background.

Emission events were classified, employing a Hidden-Markov-Model (HMM) to estimate the time-dependent state of the fluorophore (‘off’, ‘on’ or ‘blinking’) in the detection region from the photon trace. Emission that was considered valid, meaning that it was assigned the states “on” or “blinking”, was segmented into groups of 1000 or 2000 photons (thus guaranteeing unbiased position estimation) for separate localizations, from which experimental estimates of the localization precisions were also obtained.

Classified events affected strongly by out-of-focus fluorescence could be identified by high $p_0$ values close to 0.25, which is equivalent to no intensity modulation from moving the excitation doughnut through the TCP. Clear separation between centered molecules and background was possible by monitoring the time-averaged $p_0$ value. A sharp increase in the $p_0$-value to an average of 0.25 coincided with off-switching steps. This was the case for most molecule events and the $p_0$-value was not required as an identifier for valid states here, because the HMM already classifies the event correctly. However, especially in structures with a high label density, events that were strongly affected by out-of-focus fluorescence were recorded. To discard such events (which the HMM might wrongly classify as valid states), the $p_0$-value was used as a quality criterion and all re-segmented photon bunches with $p_0 >= 0.23$ were considered as background. This procedure inherently avoided artifactual localizations distributed along the scanned grid positions. Detailed post-processing parameters are listed in Table S8.

#### Cluster-analysis of the VGlut and AMPAR datasets

For the VGlut and AMPAR cluster analysis, we did not use the automated threshold determination but followed the post-processing procedure described in (8). Detailed post-processing parameters are shown in Table S9. In case of VGlut, cluster analysis is performed over localizations based on the re-segmented photon trace. In case of AMPAR clustering, molecules are assigned by combining re-segmented localizations if they come from the same emission event or if they lie closer together than 1.5nm. The MATLAB ‘dbscan’ algorithm (10), based on (11), is employed for assignment of clusters. For VGlut, the parameters of the dbscan algorithm are set to (epsilon = 20, MinPts = 15), for AMPAR to (epsilon = 20, MinPts = 3). For VGlut, a transparent circle is drawn over the localized clusters using the MATLAB function ‘viscircles’ (12). For AMPA receptor the border of the clusters was interpolated using cubic splines (MATLAB function ‘cscvn’ (13), based on (14)).

#### MINFLUX image rendering

2D MINFLUX images were rendered by plotting a Gaussian distribution for each localization (8). Overlapping Gaussian distributions were summed-up. The color maps are not linear.

3D MINFLUX images are displayed as 3D scatter plots with a marker size of 15 nm (~3σ of the localization precision) or as isosurface renderings. For the isosurface rendering, the localization data was converted to a 3D histogram using the MATLAB function ‘histcn’ (15) and the histogram data was rendered using the MATLAB ‘isosurface’ function (16).

MINFLUX images of actin in dendrites are delineated following a two-step procedure. The first step is a nearest-neighbor filtering to reject single and isolated localizations with a mean distance of more than 150 nm to the next 20 neighbors. The nearest neighbor distances were calculated using the MATLAB ‘knnsearch’ function (17), based on (18) with the number of nearest neighbors K=20. The mean over the 20 nearest neighbor distances $\langle D_{NN} \rangle$ was calculated for each localization. Only localizations with $\langle D_{NN} \rangle < 150nm$ were selected. In the second step, the

structure was reconstructed using the MATLAB shape-reconstruction function ‘alphashape’ (19) on the filtered localizations with a custom chosen alpha-radius α, that scaled anti-proportionally to the localization density in an image.
The shape reconstruction approach is illustrated in Fig. S8 B.

##### Simulation of AMPA receptors

To simulate the 2D MINFLUX images of AMPA receptors, the positions of the CAM2 labeling site (LYS471) were extracted from the protein data base file 3KG2. Randomly distributed AMPA receptors were simulated in a field of view of 1500nm×1500 nm×100 nm with a minimum distance of 10 nm to each other. Between one and four subunits were labelled and free rotation of the receptor about the optical axis as well as up to +/-45° rotation perpendicular to the optical axis was allowed. A Gaussian distribution of localizations around the receptor subunits with a sigma of 2 nm was simulated. Nearest neighbor histograms are plotted. The results are compared to random distribution of dots without the AMPA geometry and to the measured data (see Fig. S9).

### Supporting Text

#### Focal intensity distribution in tissue

MINFLUX nanoscopy works by centering an excitation-beam minimum onto an emitter. Therefore, the shape of the excitation beam needs to be known. When imaging deep into tissue, the excitation light as well as the fluorescence interact with the tissue matter, which can influence the amplitude, phase and polarization of the photons. In MINFLUX nanoscopy, the fluorescence photon rate is measured, which is proportional to the excitation intensity. From the number of photons detected, when the excitation beam is at each position in the targeted-coordinate pattern (the scan-pattern) respectively, we can estimate the position of the molecule. To get an indication on how reliable MINFLUX can work in tissue, the intensity distribution of the excitation light in tissue (point spread function (PSF)) needs to be known.

###### Modeling of depth-induced aberrations by Zernike polynomials

Assuming that the tissue slice is a homogenous optical layer with a certain isotropic refractive index and a flat surface, it only influences the phase of the photons. In this simplified model, we can calculate the PSF shape in different depths in tissue.
We compare the situation when imaging directly on the coverslip to the situation when imaging in tissue by moving the sample towards the objective by a certain distance Δh.

In the ray optical image, the light rays that come from the objective travel through the immersion medium with refractive index $n_1$ and hit the coverslip with an angle α that is dependent on the radial position ρ of the rays in the back aperture. From Snell's law, we can calculate the angle γ in the tissue sample with refractive index $n_3$:

$$\gamma = \mathrm{asin}(n_1/n_3 \cdot \sin(\alpha)). \quad (1)$$

The angle α is given by radial ray starting position $\rho$ in the objective back aperture and the focal length $f$ of the objective

$$\alpha = \tan^{-1}\left(\frac{\rho}{f}\right). \quad (2)$$

When moving the objective lens the distance Δh through the immersion oil towards the sample or increasing the thickness of the coverslip by $\Delta cs$, the ray crossing point shifts the distance $\Delta h_{\mathrm{optical}}$ into the sample:

$$\Delta h_{\mathrm{optical}} = \frac{\tan(\alpha)}{\tan(\gamma)} \cdot \Delta h + \frac{\tan(\beta)}{\tan(\gamma)} \cdot \Delta cs. \quad (3)$$

Since we use coverslips with constant thickness, we set $\Delta cs = 0$.

In the case of central rays or no refractive index mismatch between sample and immersion oil, $\alpha \approx \gamma$ and the optical depth is equal to the mechanical depth. However, this is not the case for the outer rays if $n_1 \neq n_3$, which leads to broadening of the focus, because rays with different angles α constructively interfere at different depths in the sample. Therefore, the effect of focusing a distance Δh into the sample is equivalent to a phase shift $\Delta\varphi$ in the objective back aperture:

$$\Delta\varphi(\rho, \Delta h) = \frac{2\pi}{\lambda}\left(n_1\sqrt{\rho^2 + \Delta h^2} - n_3\sqrt{\rho^2 + \Delta h_{\mathrm{optical}}^2(\rho, \Delta h)}\right). \quad (4)$$

With equations (1), (2) and (3), equation (4) can be simplified to:

$$\Delta\varphi(\rho, \Delta h) = \frac{2\pi}{\lambda}\Delta h\left(n_1 - \frac{{n_3}^2}{n_1}\right)\sqrt{1 + \frac{\rho^2}{f^2}} \ . \quad (5)$$

To get a more intuitive understanding of the influence of this phase shift onto the PSF of the microscope, we want to expand equation (5) into Zernike polynomials $Z_n^m(r, \theta)$ in cylindrical coordinates (20). Zernike polynomials are an orthogonal and complete set of functions defined on the unit circle and are commonly interpreted as optical aberrations.

Since the function $\Delta\varphi$ is radially symmetric, only the Zernike polynomials with angular index m = 0 need to be considered. These are the Zernike polynomials for piston, defocus, primary and higher order spherical aberration.

We define $\rho = r\,\rho_{max}$, with $0 \leq r \leq 1$, and rewrite equation (5) to:

$$\Delta\varphi(\mathrm{r},\,\Delta h) \;=\frac{2\pi}{\lambda}\Delta\mathrm{h}(n_1-\frac{{n_3}^2}{n_1})\sqrt{1+r^2\frac{\rho_{max}^2}{f^2}}. \tag{6}$$

As the Zernike polynomials are orthogonal and complete, we can then calculate the Zernike coefficients $c_n^{0\prime}$ for $n \;=\; 0,2,4,6,8,\dots$ via:

$$c_n^{0\prime} \;=\; 2\int_0^1 \Delta\varphi(\mathrm{r},\,\Delta h)\,Z_n^0(r)\,r\,dr. \tag{7}$$

The Zernike expansion can be done analytically, but it leads to rather lengthy terms.

Therefore, we just show the results for the numerical values $\mathrm{f} = 1.8\,\mathrm{mm}$ and $\rho_{max} = 2.43\,\mathrm{mm}$:

$$\Delta\varphi(\mathrm{r},\,\Delta h) \;\approx \tfrac{2\pi}{\lambda}\Delta\mathrm{h}(n_1-\tfrac{{n_3}^2}{n_1})$$
$$\Big(1.3688\,Z_0^0(r)+0.1938Z_2^0(r)-0.0125\,Z_4^0(r)+0.0016\,Z_6^0(r)\;-0.0003\,Z_8^0(r)\Big). \tag{8}$$

For comparison and validation of this approach, the “ground truth” phase function equation (6) is compared to its Zernike expansion up to order n = 8, equation (8).

$$\Delta\varphi(\mathrm{r},\,\Delta h) \;\approx \tfrac{2\pi}{\lambda}\Delta\mathrm{h}(n_1-\tfrac{{n_3}^2}{n_1})$$
$$\Big(1.3688\,Z_0^0(r)+0.1938Z_2^0(r)-0.0125\,Z_4^0(r)+0.0016\,Z_6^0(r)\;-0.0003\,Z_8^0(r)\Big). \tag{8}$$

A reconstruction using only piston, defocus and primary spherical aberration (n <= 4) gives agreement between the phase function and its Zernike expansion.

While piston, being only a constant phase offset, does not affect the PSF and the defocus only shifts the position of the PSF along the optical axis, the spherical aberration affects the shape of the PSF. The higher order spherical aberrations contribute much less to the phase function than the primary spherical aberration and therefore, we neglect them in the following. We measure the excitation PSF by scanning the excitation beam over a point-like object (i.e. a fluorescent bead or a small caveolin1-cluster). We can further determine the Zernike coefficient for spherical aberration by adjusting the Zernike coefficient for the phase correction of the excitation beam via a SLM until the PSFs no longer show the aberration linked to the respective Zernike polynomial. From the equation (8), we get for the spherical aberration Zernike coefficient in the objective back aperture:

$$c_4^{0\prime} \;=\; -0.0125\,\frac{2\pi}{\lambda}\,\Delta\mathrm{h}\left(n_1-\frac{n_3^2}{n_1}\right). \tag{9}$$

To compare with the experimental data, we normalize the Zernike coefficient $c_4^{0\prime}$ to k = $\frac{2\pi}{\lambda}$ and correct with a pupil magnification factor $m_p \;=\; (\frac{4}{3})^4$ since the SLM is imaged with a 4f-system with a focal length ratio of 3:4 onto the objective back aperture (the scaling factor is determined as described in (21)). We further need to multiply with a phase scaling factor of $m_r = 2$ , since the SLM is used in reflection mode and therefore the phase correction is applied twice. Therefore, equation (9) is rewritten as:

$$c_4^0 = -0.025 \ (\frac{4}{3})^4 \ \frac{2\pi}{\lambda} \Delta h(n_1 - \frac{{n_3}^2}{n_1}), \quad (10)$$

resulting in an equation that links the spherical aberration to the refractive index of the sample.

**Calibrating refractive index measurements by spherical aberration determination**

First, we measured the spherical aberrations in agarose-sucrose pads with embedded fluorescent beads. We determined the Zernike coefficient $c_4^0$ in different depths in the sample. The agarose-sucrose mixture allows us to tune the refractive index with the content of sucrose, similar to how it is described in (4). We measured the refractive index of the resulting sample using a critical – angle dispersion refractometer (Schmidt + Haensch, ATR-L; accuracy: 0.0005 RI at 20°C). Since the agarose-sucrose pad is homogenous as well as transparent in the optical wavelength range and does not affect the polarization of the light, it is a good model system for testing the influence of the refractive index mismatch on the focal light intensity distribution. We used a silicon oil immersion objective for the PSF measurements. Exemplary images of measured PSFs in 0, 40 and 80 µm imaging depths are shown in Fig. S4G-I. We determined the Zernike coefficient $c_4^0$ experimentally by adding a Zernike-based phase correction to the SLM until the aberrations visible in the PSF measurements with different phase masks are minimal. We fit a polynomial of first order to the spherical aberration coefficient over the depth. This is shown in Fig. S4A-C. From the gradient of the linear fit $m = \frac{c_4^0}{\Delta h}$, we can then calculate the refractive index of the sample by solving equation (10) for $n_3$:

$$n_3 \approx -0.177 \sqrt{-405\, m + 32\, n_1^2}. \quad (11)$$

The refractive index of the silicon oil immersion is $n_1 = 1.406$ at RT (as specified by the manufacturer). The refractive indices calculated from the PSF measurements with equation (11)for the samples shown in Fig. S4 A-C are shown in Table S1. They match well with the refractive indices measured by the refractometer. This can be seen as a validation of our theoretical model of the phase difference due to refractive index mismatch in equation (4). Similar aberration measurements in a gel sample were demonstrated in (22), also showing a linear dependence of spherical aberration on depth.

**Measurement of PSFs in tissue samples**

Next, we wanted to investigate aberrations in biological tissue samples. To this end, we measured PSFs of the beads injected into the fixed mouse brain tissue. In addition, we investigated the PSFs of small (<= 100nm) spherical agglomerates of caveolin-1 labelled with AF647 in similar brain tissue sample. All measurements were performed in LI of visual cortex. The results are shown in Fig. S4 J-L and D-F. Noticeably, the PSFs in the tissue samples are more elongated along the optical axis and the zero of the z-doughnut PSF is filled up more in the depth comparing to the ones in the agarose-sucrose sample. This can be accounted for by the non-negligible absorption and scattering in the tissue. The influence of absorption and scattering of the tissue on the light intensity (I) can be described by the Lambert-Beer law:

$$I = I_0 \, e^{-\mu \, x}, \qquad (12)$$

with the extinction coefficient $\mu = \mu_a + \mu_s$ composed of the absorption and scattering coefficients and x being the optical path length in the tissue. $1/\mu$ then describes the mean free path length in tissue. From attenuation measurements in acute swine brain slices (grey matter) (23), a mean free path length of ~63 µm could be determined. This value may however not be exactly the same for our fixed mouse brain tissue slices. As the optical path length in the tissue is different for rays starting at different positions in the objective back aperture, the outer rays are far more attenuated than the central ones, explaining qualitatively the observed elongation along the optical axis in the depth of the tissue as well as the filling up of the z-doughnut zero.

Still, like in the agarose-sucrose sample, the zero of the vortex PSF is quite robust in the depth, which is the most important for MINFLUX localizations. The elongation of the PSF along the optical axis may, however, increase background from the emitters located in different z-planes. As shown in (24), spherical aberrations do not induce bias in 2D MINFLUX, even if the actual focal intensity distribution is not measured and accounted for in the position estimation.

For minimizing the effects of the phase difference onto the PSF, good refractive index matching of the sample to the objective and immersion medium is crucial. Due to the high absorption and scattering of tissue, it is not possible to directly measure its refractive index with the refractometer. We can however, deduce the refractive index from the spherical aberration measurements using equation $(11)$. The resulting refractive indices calculated for the tissue samples are shown in Fig. S4 M. The mean value of calculated refractive index of tissue is about ~1.377, which is in agreement with OCT measurements of the refractive index in living brain tissue of rats (25) and mice (26).

As shown in Fig. S4 M, matching of refractive indices between immersion oil and sample minimizes the gradient of the primary spherical aberration with depth. Compared to water ($n_1 = 1.33$), glycerol ($n_1 = 1.456$) and Leica type F immersion oil ($n_1 = 1.518$), silicon immersion oil with

$n_1 = 1.406$ matches best the refractive index of tissue. Mixtures of immersion liquid like glycerol and water would match the refractive index of tissue better. However, the high evaporative rates of the components of such mixtures lead to refractive index changes over time, making them less advantageous for MINFLUX nanoscopy, because the duration of MINFLUX acquisitions can be up to several hours. As the silicon oil is resistant to evaporation, helps to mitigate refractive index mismatch and reduces spherical aberrations, we decided to use a silicon oil immersed objective for MINFLUX nanoscopy at several tens of micrometer depth in biological tissue.

The problem of absorption and scattering of tissue may be reduced by clearing of the tissue, but this has the disadvantage of more complicated sample preparation and possible sample preparation artifacts.
We reason, based on our study of the focal intensity distribution in tissue, that 2D MINFLUX relying on the vortex excitation beam should be possible up to ~80µm deep in tissue even without active aberration correction, when using a silicon oil objective to match the tissue refractive index.

### Figures S1 to S11

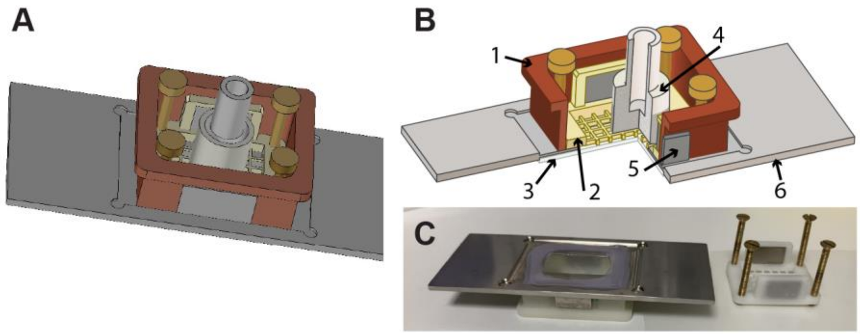


**Fig. S1: Chamber for living slice imaging. A, B** Schematic drawing of chamber with oxygen supply: **1**-basin, **2**-grid, **3**-coverslip, **4**-percolator delivering gas, **5**-magnet, **6**-steel slide. **C** Life imaging chamber mounted on slide (left) and the grid for fixation of the tissue slice (right).

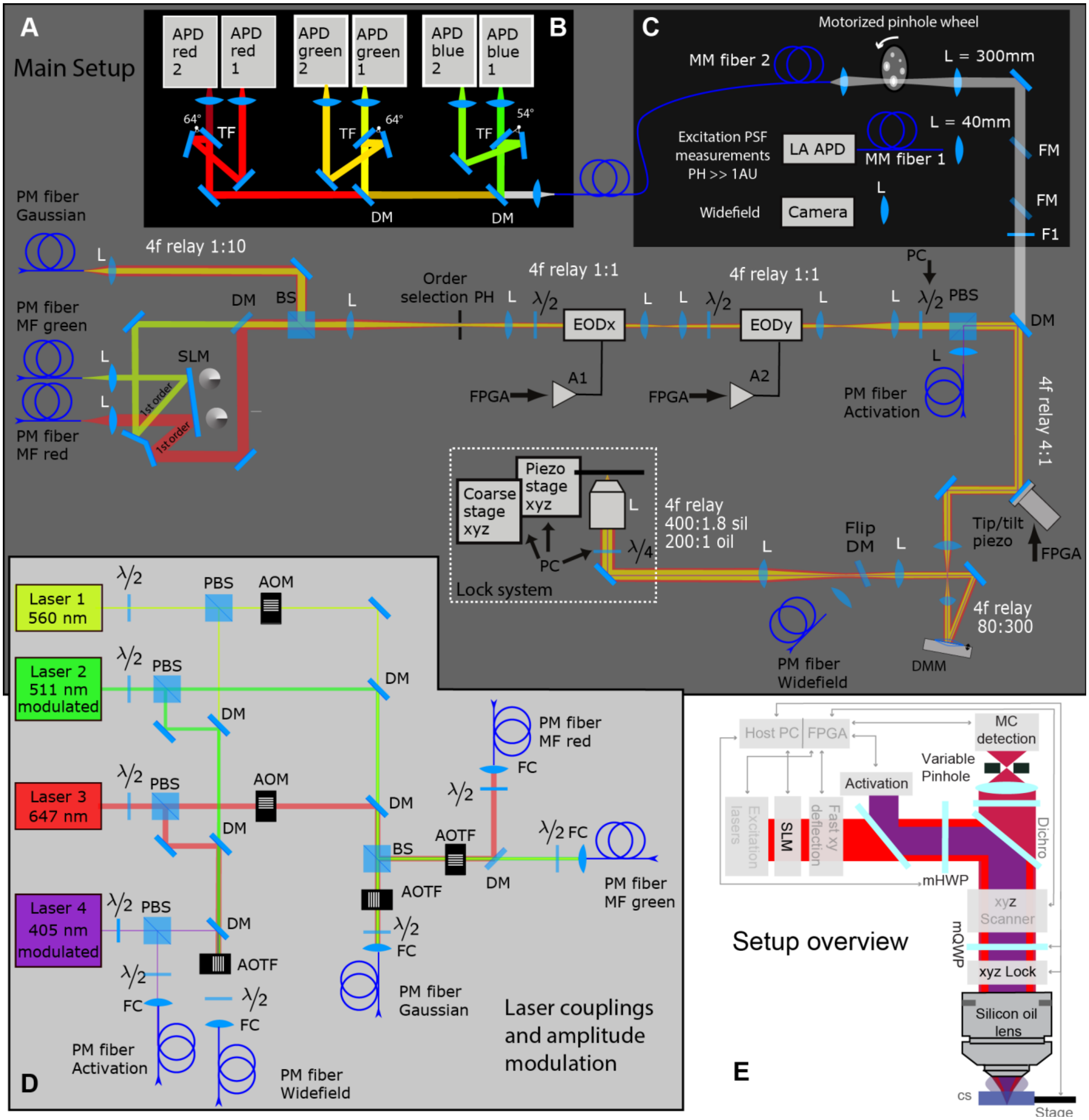


**Fig. S2: Overview of MINFLUX setup**. Detailed drawing of setup with all crucial components (not to scale). **A** Main setup consisting of the excitation, activation and detection beam paths, phase modulation, scanner system, microscope stage and objective. Depth adaptable lock system shown in detail in Fig. 2. **B** Light-proof multicolor detection box for spectrally separated confocal detection. **C** Detection box with the Camera for widefield imaging, the Large Area (LA) APD for PSF measurements and the variable confocal pinhole. Widefield camera and LA APD not illuminated. **D** Excitation and activation laser box with power modulator. Boxes are linked by optical fibers for better stability and exterior light protection. **E** MINFLUX setup overview from Fig. 1A shown again as an overview. Abbreviations for components explained in Table S2.

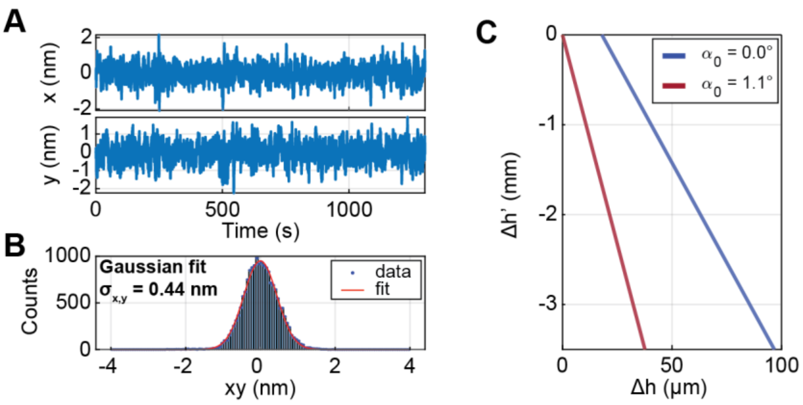


**Fig. S3: Depth-adaptable focus lock system. A** Exemplary time traces of the focus lock position deviations in x and y for a stability measurement directly on the coverslip. Measurement was performed in an agarose-sucrose gel with refractive index of 1.38. **B** A histogram of combined projection of x and y focus lock position deviations. Gaussian fit to the data shown in red. Resulting focus lock precision for a measurement of ~10 min is 0.44nm. **C** Simulation of beam movement on camera $\Delta h'$ due to sample movement along the optical axis $\Delta h$ for different angles $\alpha_0$ of the incoming beam that can be selected by the absolute positionable mirror.; $\sigma_{xy}$ – position stability.

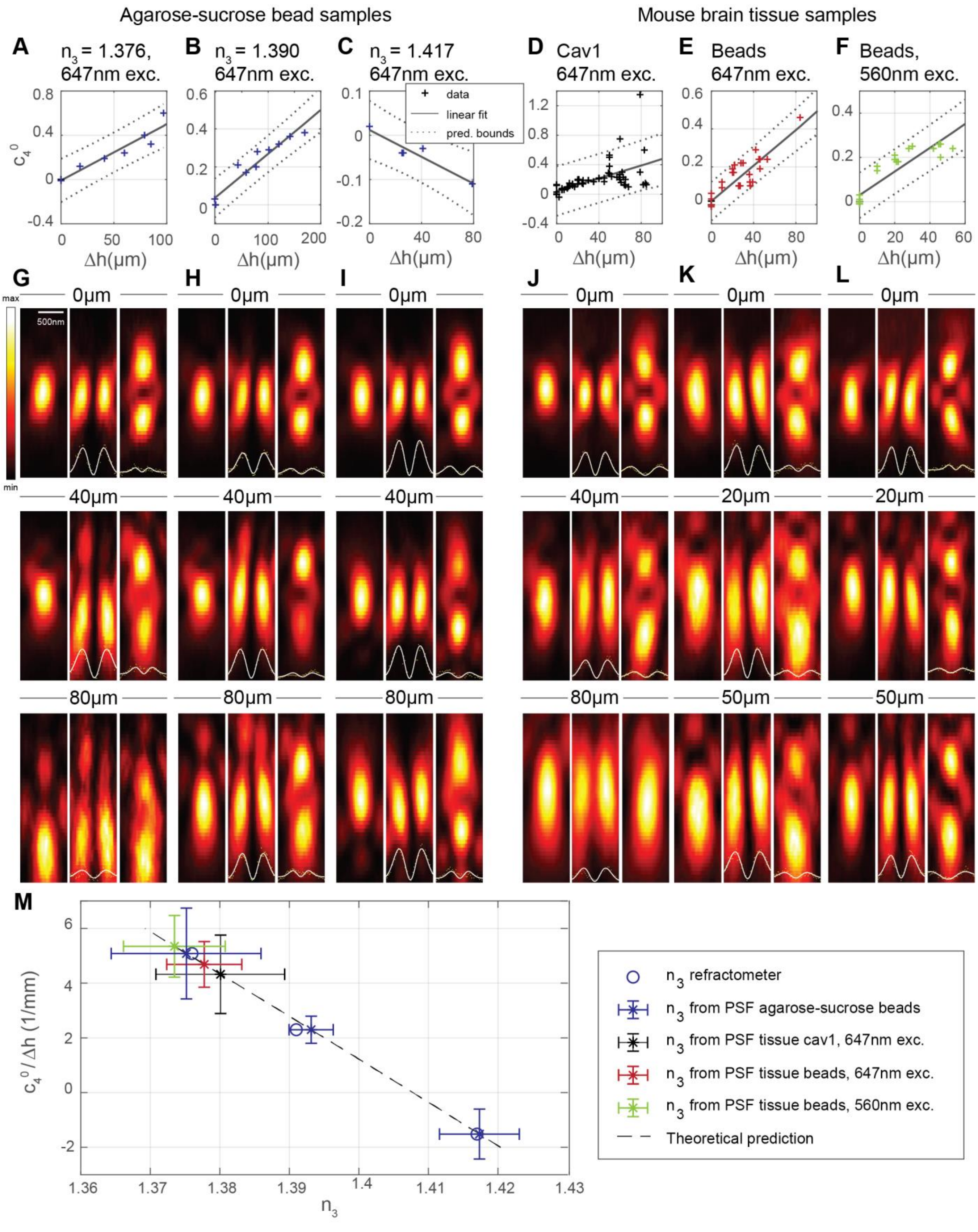

Agarose-sucrose bead samples
Mouse brain tissue samples
A $n_3$ = 1.376, 647nm exc.
B $n_3$ = 1.390 647nm exc.
C $n_3$ = 1.417 647nm exc.
D Cav1 647nm exc.
E Beads 647nm exc.
F Beads, 560nm exc.
data
linear fit
pred. bounds
$c_4^0$
Δh(μm)
G
H
I
J
K
L
0μm
40μm
20μm
50μm
80μm
500nm
max
min
M
$c_4^0$ / Δh (1/mm)
$n_3$
$n_3$ refractometer
$n_3$ from PSF agarose-sucrose beads
$n_3$ from PSF tissue cav1, 647nm exc.
$n_3$ from PSF tissue beads, 647nm exc.
$n_3$ from PSF tissue beads, 560nm exc.
Theoretical prediction

**Fig. S4: Focal intensity distributions (PSFs) in different imaging depths. A**-**F** Spherical aberration over depth measured in agarose-sucrose samples (A-C) and in mouse brain tissue sample (D-F) for 647nm (A-E) or 560nm (F) excitation wavelength. **G**-**L** Axial cuts through exemplary PSFs for three different imaging depths for corresponding sample type. Imaging depths were rounded to 10 µm. Regular focus, doughnut and z-doughnut PSFs are shown. Line profiles perpendicular to the optical axis characterizing the zero are shown together with a doughnut shaped fit for each doughnut PSF and z-doughnut PSF. While the zero contrast of the doughnut is still good in the depth, the zero contrast of the z-doughnut decreases rapidly. **M** Data summary showing the relation between the depth-dependent spherical aberration and the refractive index of the sample together with the result of the refractometer measurements of the agarose-sucrose gels. cav1 – caveolin-1; $\Delta h$ - depth; $n_3$ - refractive index; exc. - excitation; pred. – prediction.

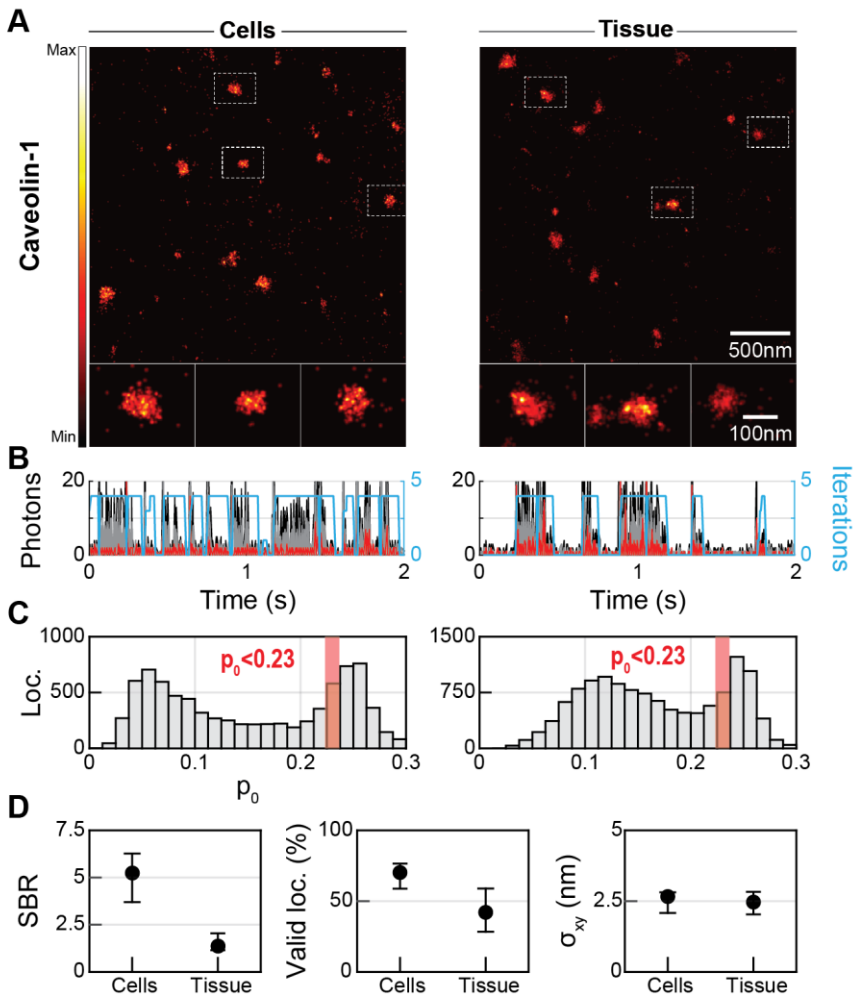


**Fig. S5: MINFLUX imaging of Caveolin1 in cells and tissue. A** Overview MINFLUX images of Caveolin-1 in cultured U-2 OS cells and mouse brain tissue close to the coverslip (upper panel). Lower panel shows zoom-in pictures of three selected Caveolin1-clusters found in delineated areas indicated in overviews. **B** Selected photon trace showing molecule emission events and iterative centering of the targeted-coordinate pattern onto the molecules for measurement shown in A. **C** Histograms of p0-values of localizations in the images from A. **D** Median values and interquartile range of SBR, ratio of valid localizations and localization precision calculated for MINFLUX images of caveolin1 in cells (6 images, 22478664 localizations) and tissue (23 images, 703924 localizations). Loc. - localizations; SBR - signal to background ratio; $\sigma_{xy}$ - localization precision.

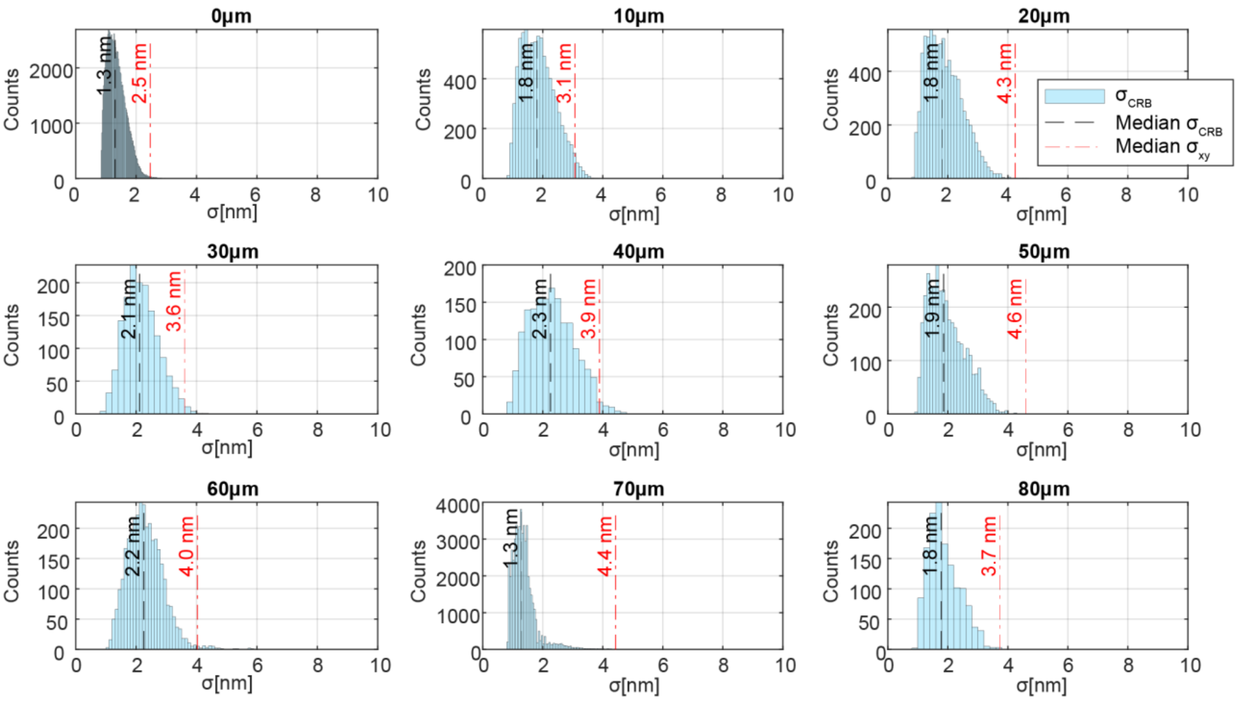


**Fig. S6: MINFLUX imaging of Caveolin1 in different depths of mouse brain tissue.** Cramér-Rao-bound ($\sigma_{CRB}$) calculated from experimentally determined signal-to-background ratio (SBR) compared to the measured localization precision ($\sigma_{xy}$). SBR - signal to background ratio; σ - localization precision.

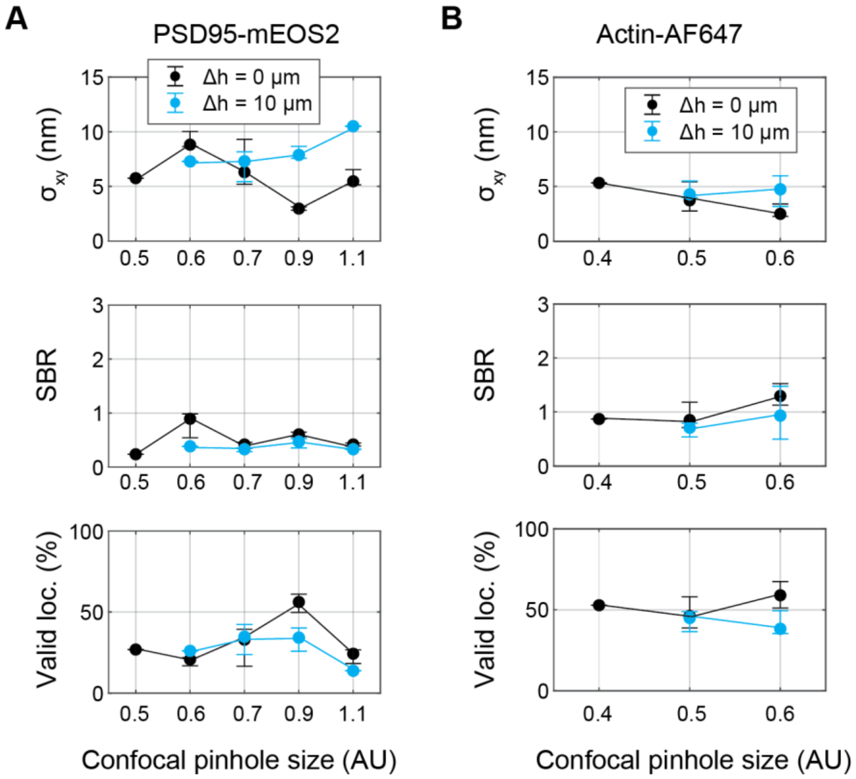


**Fig. S7: Influence of confocal pinhole size on the MINFLUX metrics.** Localization precision, SBR and percentage of valid localizations were evaluated for different confocal pinhole sizes directly on the cover slip (0 µm) and at ~10 µm depth for 2 different samples; PSD95-mEOS2 (**A**) and Actin-Alexa Fluor 647 (**B**). Median values with interquartile range are plotted. Sample size for PSD95 with 0.5 AU: 1 image, 23729 localizations, PSD95 with 0.6 AU: 4 images, 2071616 localizations, PSD95 with 0.7 AU: 42 images, 1847111 localizations:PSD95 with 0.9 AU: 27 images, 335680 localizations, PSD95 with 1.1 AU: 8 images, 448405 localizations. AF647 with 0.4 AU: 1 image, 5765 localizations, AF647 with 0.5 AU: 64 images, 37515021 localizations, AF647 with 0.6 AU: 24 images, 16798087 localizations. $\Delta h$ - depth; AF - Alexa Fluor; loc. - localizations; SBR - signal to background ratio; $\sigma_{xy}$ - localization precision.

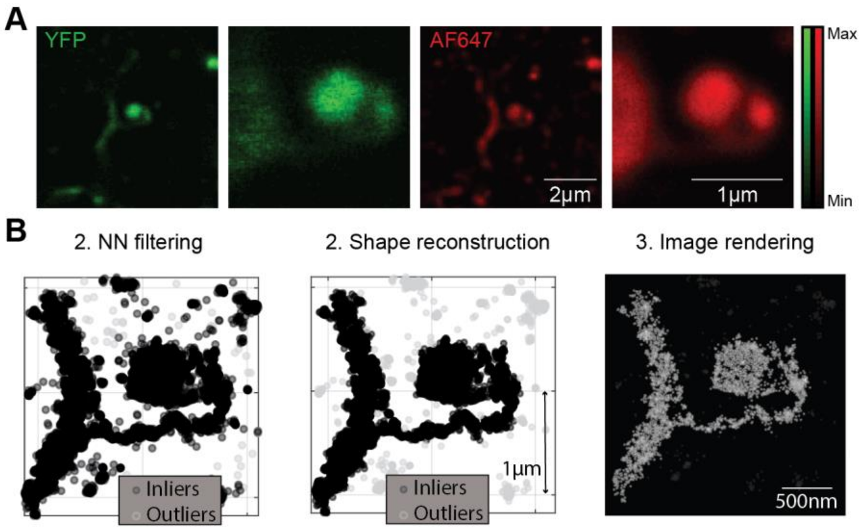


**Fig. S8: Delineation of actin in dendritic spines. A** confocal overview images and zoom-ins to the region of interest shown for YFP and Alexa Fluor 647 channel. **B** Raw MINFLUX localizations from the ROI with nearest neighbor filtering (NN filtering, left image), shape reconstruction (center image) and image rendering as sum of Gaussian distributions (right image).

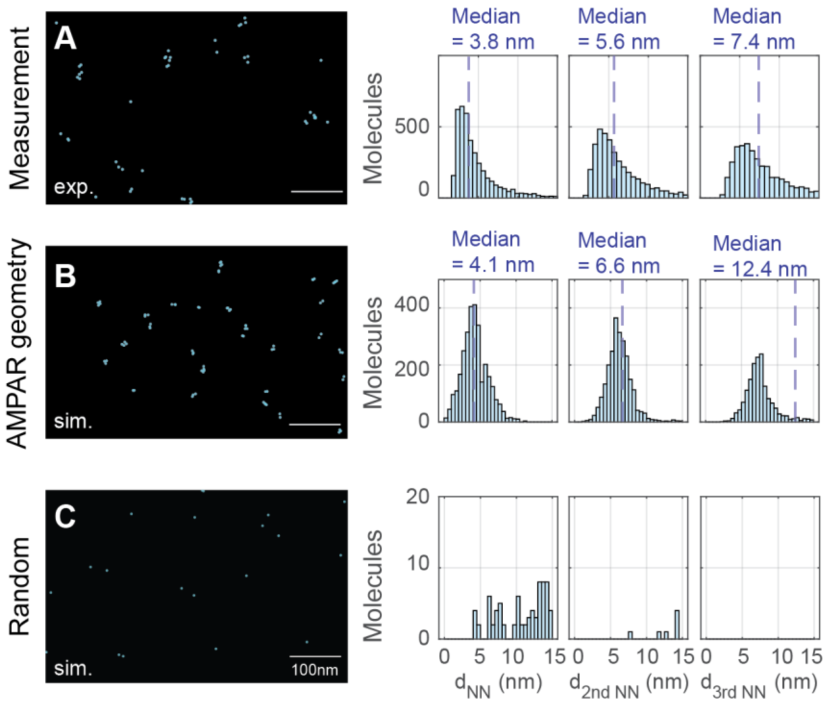


**Fig. S9: Comparison of AMPA receptor measurement with simulation. A** Selected field of view of 2D MINFLUX image. On the right, nearest neighbor distances, second nearest neighbor distances and third nearest neighbor distances between AMPA subunit localizations are shown. Localizations with distances below 1.5 nm are assigned to the same molecule. Nearest neighbor distances are calculated only for clusters with at least two subunits within 15 nm. **B** Selected field of view of simulation of AMPA receptors relying on the protein data base AMPA label site geometry. Nearest neighbor distances between simulated subunits. **C** Same for simulation of random localizations. Exp. – experimental data, sim. – simulated data.

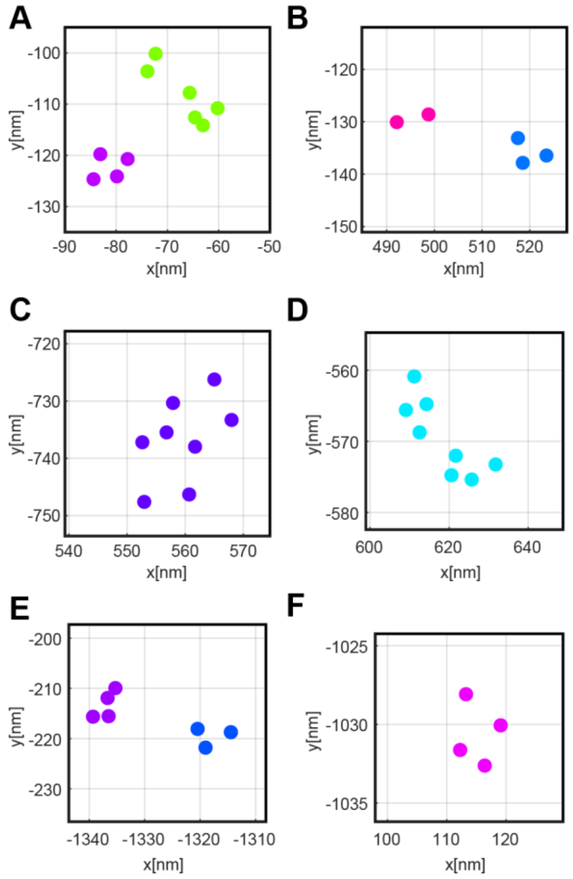


**Fig. S10: AMPAR geometries overview.** Selected ROIs (A-F) from the AMPA receptor image shown in Fig. 5E. Colors are assigned by DBSCAN cluster algorithm.

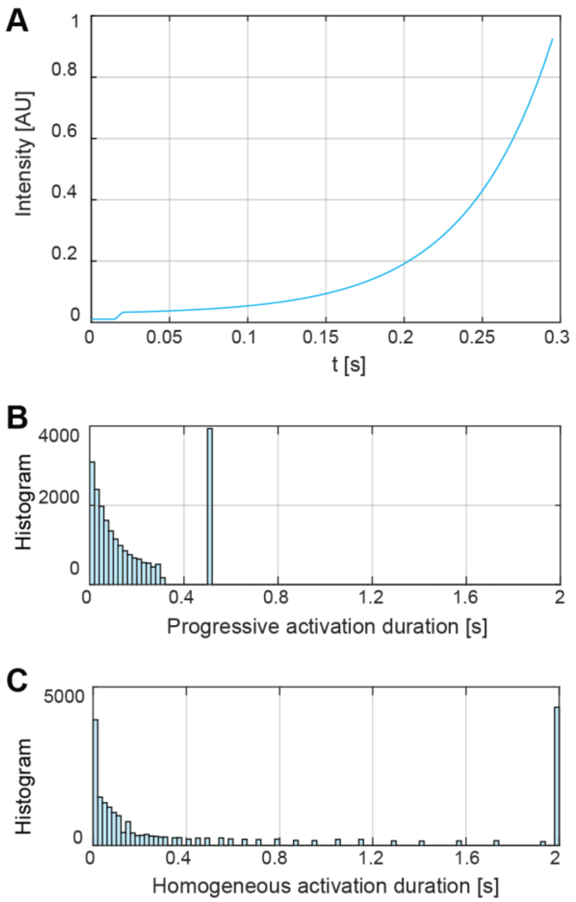


**Figure S11: Progressive Activation. A** Temporal change of activation intensity. **B** A histogram of recorded activation durations during an image acquisition. The large peak at 0.5 s are the instances where no activation of a molecule happened (including the time for the piezo mirror jump of 200 ms). **C** Estimated histogram of activation durations, based on the doses per activation in **B** and a homogeneous activation intensity of 2 % of the maximum intensity in **B**.

### Tables S1 to S10

Table S1: Refractive indices of agarose-sucrose samples calculated from measurements of primary spherical aberrations compared to the refractometer measurements.

| Sample (shown in Fig. S2) | A | B | C |
|---|---|---|---|
| Concentration (m/v) of agarose (%) | 0.8 | 1.98 | 2.37 |
| Concentration (m/v) of sucrose (%) | 33.4 | 31.8 | 68.0 |
| $n_3$ (refractometer at 656.3 nm) | 1.376 | 1.391 | 1.417 |
| $n_3$ (aberration measurements) | 1.375 | 1.393 | 1.417 |
| 95% confidence interval (from aberration measurements) | [1.364,1.386] | [1.393,1.396] | [1.412,1.423] |

Table S2: List of component abbreviations used in the detailed setup overview in Fig. S2.

| Component | Abbreviation |
|---|---|
| Half-wave plate (HWP) | $\lambda/2$ |
| Quarter-wave plate (QWP) | $\lambda/4$ |
| DM | Dichroic mirror |
| PBS | Polarizing beam splitter |
| BS | Beam splitter |
| AOM | Acousto-optic modulator |
| AOTF | Acousto-optic tunable filter |
| PM fiber | Polarization maintaining optical fiber (single mode) |
| FC | Fiber collimator/coupler |
| TF | Tunable filter |
| APD | Avalanche photodiode |
| MM fiber | Multimode optical fiber |
| EOD | Electro optical deflectors |
| SLM | Spatial light modulator (blazed grating to reflect into first order, phase masks) |
| L | Lens |
| FPGA | Field programmable gate array |
| PH | Pinhole |
| PC | Computer |
| Flip DM | Flip dichroic mirror (flip into beam path for widefield imaging) |
| DMM | Deformable membrane mirror |
| FM | Flip mirror |
| F1 | Notch filters (Laser wavelengths, focus lock wavelengths) |

Table S3: Localization patterns for 2D MINFLUX adapted from (8).

| Excitation wavelength (nm) | Iteration Index | Beam shape | TCP | L (nm) | Max. #Phot. to | Est. param. |
|---|---|---|---|---|---|---|
| 560 | 1 | regular focus | 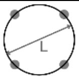 | 300 | 100 | 0.8 |
| | 2 | regular focus | 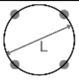 | 300 | 100 | 0.8 |
| | 3 | doughnut | 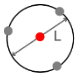 | 150 | 100 | 0.883, 6.623 |
| | 4 | doughnut | 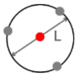 | 100 | 250 | 0.728, 9.15 |
| | 5 | doughnut | 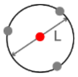 | 100 | 10000 | 0.729, 9.114 |
| 647 | 1 | regular focus | 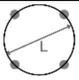 | 300 | 100 | 0.8 |
| | 2 | regular focus | 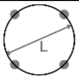 | 300 | 100 | 0.8 |
| | 3 | doughnut | 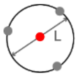 | 150 | 100 | 0.875, 6.857 |
| | 4 | doughnut | 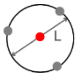 | 100 | 250 | 0.719, 9.305 |
| | 5 | doughnut | 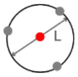 | 100 | 10000 | 0.719, 9.269 |

Table S4: Overview of 2D MINFLUX measurement parameters for different samples. Laser powers are measured close the objective. Localization patterns are detailed in Table S3.

| Sample | U-2 OS cells αCaveolin1-αAF647 | Tissue αCaveolin1-αAF647 | Tissue LifeAct - EYFP αGFP - αAlexa Fluor 647 | Tissue AMPAR - CAM2 Alexa Fluor 647 | Tissue PSD95-mEOS2 |
|---|---|---|---|---|---|
| Excitation wavelength (nm) | 647 | 647 | 647 | 647 | 560 |
| Laser power for excitation with doughnut focus (µW) | 100 | 100 | 100 | 100 | 14-40 |
| Laser power for excitation with regular focus (µW) | 60 | 60-80 | 60 | 60 | 8-25 |
| Laser power for activation @ 405nm (µW) | Progressive activation up to 1 | Progressive activation up to 0.1- 1.4 | Progressive activation up to 1 | Progressive activation up to 0.1 | 0.006-1.1 |
| Confocal pinhole diameter (AU) | 0.62 | 0.46 -0.54 | 0.46-0.62 | 0.62 | 0.53-0.9 |
| Pixel pitch (distance between mosaic positions at which we activate) (nm) | 200 | 200 | 200 | 200 | 150-200 |
| Maximum activation time at mosaic index (s) | 1.2 | 0.3-1.2 | 0.3-1.2 | 0.6 | 0.3-1.2 |
| Reset of activation time with molecule | yes | yes | yes | yes | yes |
| Maximum time at mosaic position (s) | 10 | 10-20 | 10 | 10 | 10-15 |
| Localization pattern | 2D MINFLUX 647nm excitation | 2D MINFLUX 647nm excitation | 2D MINFLUX 647nm excitation | 2D MINFLUX 647nm excitation | 2D MINFLUX 560nm excitation |

Table S5: 3D MINFLUX localization schemes for z-doughnut vs. doughnut-regular focus comparison measurements and 3D two-color measurements. z-doughnut localization scheme adapted from (8).

| Main excitation beam(s) | Localization pattern | | | | | | |
|---|---|---|---|---|---|---|---|
| Doughnut 647nm + regular focus 647nm | Iter. Ind. | Beam shape | TCP | # exposures | L (nm) | Max. #Phot. to collect | Est. param. |
| | 1 | regular focus | 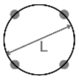 | 4 | 300 | 100 | 0.8 |
| | 2 | Regular focus | 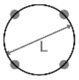 | 4 | 300 | 100 | 0.8 |
| | 3 | Regular focus | 1D, z | 2 | 1200 | 100 | 0.15 |
| | 4 | regular focus | 1D, z | 2 | 1200 | 100 | 0.15 |
| | 5 | regular focus | 1D, z | 2 | 1200 | 100 | 0.15 |
| | 6 | doughnut | 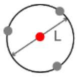 | 4 | 150 | 100 | 0.89, 7.18 |
| | 7 | doughnut | 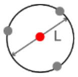 | 4 | 100 | 200 | 0.57, 10.8 |
| | 8 | doughnut/ regular focus | 3D | 6 | 100, 750 | 10000 | 0.57, 10.8, 0.35 |

z-doughnut 647nm

| Iter. Ind. | Beam shape | TCP | # exposures | L (nm) | Max. #Phot. to | Est. param. |
|---|---|---|---|---|---|---|
| 1 | regular focus | 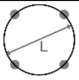 | 4 | 300 | 100 | 0.8 |
| 2 | regular focus | 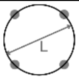 | 4 | 300 | 100 | 0.8 |
| 3 | regular focus | 1D, z | 2 | 1200 | 100 | 0.15 |
| 4 | z-doughnut | 1D, z | 2 | 300 | 100 | 0.55 |
| 5 | z-doughnut | 1D, z | 2 | 200 | 150 | 0.55 |
| 6 | z-doughnut | 3D | 7 | 150 | 150 | 0.88, 23.5 |
| 7 | z-doughnut | 3D | 7 | 100 | 200 | 0.58, 31.5 |
| 8 | z-doughnut | 3D | 7 | 100 | 10000 | 0.58, 31.3 |

| Two-color measurements with doughnut 560nm | Iter. Ind. | Beam shape | TCP | # exposures | L (nm) | Max. #Phot. to collect | Est. param. |
|---|---|---|---|---|---|---|---|
| | 1 | regular focus | 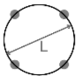 | 4 | 300 | 100 | 0.8 |
| | 2 | regular focus | 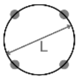 | 4 | 300 | 100 | 0.8 |
| | 3 | regular focus | 1D, z | 2 | 1200 | 100 | 0.5 |
| | 4 | regular focus | 1D, z | 2 | 1200 | 100 | 0.5 |
| | 5 | regular focus | 1D, z | 2 | 1200 | 100 | 0.5 |
| | 6 | doughnut | 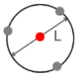 | 4 | 150 | 100 | 0.89, 7.18 |
| | 7 | doughnut | 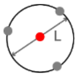 | 4 | 100 | 200 | 0.57, 10.8 |
| | 8 | doughnut/ regular focus | 3D | 6 | 100, 750 | 10000 | 0.57, 10.8, 0.35 |

| | Iter. Ind. | Beam shape | TCP | # exposures | L (nm) | Max. #Phot. to collect | Est. param. |
|---|---|---|---|---|---|---|---|
| Two-color measurements with doughnut 647nm | 1 | regular focus | 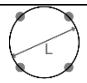 | 4 | 300 | 100 | 0.8 |
| | 2 | regular focus | 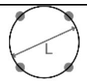 | 4 | 300 | 100 | 0.8 |
| | 3 | regular focus | 1D, z | 2 | 1200 | 100 | 0.5 |
| | 4 | regular focus | 1D, z | 2 | 1200 | 100 | 0.5 |
| | 5 | regular focus | 1D, z | 2 | 1200 | 100 | 0.5 |
| | 6 | doughnut | 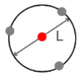 | 4 | 150 | 100 | 0.89, 7.18 |
| | 7 | doughnut | 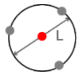 | 4 | 100 | 200 | 0.57, 10.8 |
| | 8 | doughnut/ regular focus | 3D | 6 | 100, 750 | 10000 | 0.57, 10.8, 0.35 |

Table S6: 3D MINFLUX acquisition parameters for comparison measurements between z-doughnut and doughnut/regular focus. Localization schemes are detailed in Table S5.

| Sample | Tissue LifeAct-EYFP antiGFP -antiAF647 | Tissue LifeAct-EYFP antiGFP -antiAF647 |
|---|---|---|
| Excitaion wavelength (nm) | 647 | 647 |
| Laser power for excitation with doughnut focus (µW) | 90 | 90 |
| Laser power for excitation with regular focus focus (µW) | 90 | 90 |
| Laser power for activation @ 405nm (µW) | Progressive activation up to 0.1 | Progressive activation up to 0.1 |
| Confocal pinhole diameter (AU) | 0.6 | 0.6 |
| Pixel pitch (distance between mosaic positions at which we activate) (nm) | 200 | 200 |
| Maximum activation time at mosaic index (s) | 0.6 | 0.6 |
| Reset of activation time with molecule | yes | yes |
| Maximum time at mosaic position (s) | 10 | 10 |
| Localization pattern | doughnut/regular focus | z-doughnut |

Table S7: 3D 2-color MINFLUX acquisition parameters. Localization schemes are detailed in Table S5.

| Sample | Tissue PSD95-mEOS2 LifeAct-EYFP antiGFP-antiAF647 | Tissue PSD95-mEOS2 LifeAct -EYFP antiGFP-antiAF647 | Tissue mEOS2-PSD95 AMPA CAM2-AF647 (live stained) | Tissue mEOS2-PSD95 AMPA CAM2-AF647 (live stained) |
|---|---|---|---|---|
| Excitation wavelength (nm) | 647 | 560 | 647 | 560 |
| Laser power for excitation with doughnut focus (µW) | 100 | 25 | 100 | 25 |
| Laser power for excitation with regular focus (µW) | 80 | 25 | 80 | 25 |
| Laser power for activation @ 405nm (µW) | (progressive activation to) 0.05-0.1 | (progressive activation to) 0.006-0.1 | 0.1 | 0.1 |
| Confocal pinhole diameter (AU) | 0.6 | 0.6 | 0.6 | 0.6 |
| Pixel pitch (distance between mosaic positions at which we activate) (nm) | 150-200 | 150 | 150-200 | 150 |
| Maximum activation time at mosaic index (s) | 0.3 | 0.3 | 0.3 | 0.3 |
| Reset of activation time with molecule | yes | yes | yes | yes |
| Maximum time at mosaic position (s) | 10 | 10 | 10 | 10 |
| Localization pattern | doughnut/ regular focus | doughnut/ regular focus | doughnut/ regular focus | doughnut/ regular focus |

Table S8: Fluorophore-related adaptation of parameters for post processing of MINFLUX traces of the depth imaging series and 2-color images in 2D and 3D. The $p_{0,\ max}$ -value for filtering of the localizations is set in respect to the "Background peak" center $\mu_2$ of the $p_0$ histogram. For 2D MINFLUX, the value $\mu_2 = 0.25$.

| Fluorophore | Actin AF647<br>Cav1 AF647 | mEOS2 | AMPA AF647 (live stained) | mEOS2 (in live stained slice) |
|---|---|---|---|---|
| HMM parameters | $t_{On}$ = 0.5s<br>$t_{Off}$ = 0.1s<br>$t_{Blink,On}$ = 0.001s<br>$t_{Blink,Off}$ = 0.0001s | $t_{On}$ = 5s<br>$t_{Off}$ = 1s<br>$t_{Blink,On}$ = 0.005s<br>$t_{Blink,Off}$ = 0.0005s | $t_{On}$ = 0.5s<br>$t_{Off}$ = 0.1s<br>$t_{Blink,On}$ = 0.001s<br>$t_{Blink,Off}$ = 0.0001s | $t_{On}$ = 5s<br>$t_{Off}$ = 1s<br>$t_{Blink,On}$ = 0.005s<br>$t_{Blink,Off}$ = 0.0005s |
| Molecule threshold for post processing | Automatic, quantile range [0,0.95] | Automatic, quantile range [0,0.9] | Automatic, quantile range [0,0.95] | Automatic, quantile range [0,0.9] |
| Number of photons for re-segmentation | 2000 | 1000 | 2000 | 1000 |
| Minimum number of photons for localization | 50 | 100 | 50 | 100 |
| $p_{0,\ max}$ | $0.92^*\mu_2$ | $0.92^*\ \mu_2$ | $0.8^*\ \mu_2$ | $0.83^*\ \mu_2$ |

Table S9: Postprocessing parameters for the 2D AMPAR and VGlut datasets.

| Sample | AMPAR Alexa Fluor 647 | VGlut Alexa Fluor 647 |
|---|---|---|
| HMM parameters | $t_{On}$ = 0.5s<br>$t_{Off}$ = 0.1s<br>$t_{Blink,On}$ = 0.001s<br>$t_{Blink,Off}$ = 0.0001s | $t_{On}$ = 0.5s<br>$t_{Off}$ = 0.1s<br>$t_{Blink,On}$ = 0.001s<br>$t_{Blink,Off}$ = 0.0001s |
| Molecule threshold for post processing (kHz) | 20 | 20 |
| Number of photons for re-segmentation | 2000 | 2000 |
| Minimum number of photons for localization | 100 | 100 |
| $p_{0, max}$ | 0.2 (corresponds to $0.8\mu_2$) | 0.23 (corresponds to $0.92\mu_2$) |

Table S10: Number of MINFLUX images and localizations from which the statistics shown in Fig. 4C-E are calculated.

| Sample | Number of localizations | Number of images |
|---|---|---|
| Tissue, PSD95-mEOS2 | 10332325 (~0µm) | 55 (~0µm) |
| | 3374698 (~10µm) | 29 (~10µm) |
| | 6955906 (~20µm) | 10 (~20µm) |
| | 1428416 (~30µm) | 5 (~30µm) |
| | 255912 (~40µm) | 8 (~40µm) |
| | 10815 (~50µm) | 1 (~50µm) |
| Living tissue, PS95-mEOS2 | 4116582 (~0µm) | 24 (~0µm) |
| | 4380655 (~10µm) | 15 (~10µm) |
| | 17250 (~20µm) | 1 (~20µm) |
| | 16625 (~50µm) | 1 (~50µm) |
| Tissue, actin-AF647 | 108281545 (~0µm) | 38 (~0µm) |
| | 24485604 (~10µm) | 30 (~10µm) |
| | 2062425 (~20µm) | 7 (~20µm) |
| | 3786615 (~30µm) | 6 (~30µm) |
| | 4158935 (~40µm) | 7 (~40µm) |
| | 6354430 (>~50µm) | 3 (>~50µm) |

### Legends for Movies S1 to S3

Movie S1: Animation of Fig. 7A. The two-color 3D data of actin and PSD95 is rotated. Zoom-in to the post-synapse. Rotation of only PSD95 localizations.

Movie S2: Animation of Fig. 7C. The two-color 3D data of actin and PSD95 is rotated. Zoom-in to two post-synapses and rotation of the PSD95 localizations.

Movie S3: Animation of Fig. 7D. The two-color 3D data of AMPAR and PSD95 is rotated. Zoom-in to the post-synapse and rotation of the post-synapse.
