## Supplementary figures and images for "MINFLUX fluorescence nanoscopy in biological tissue"

### MovieS1

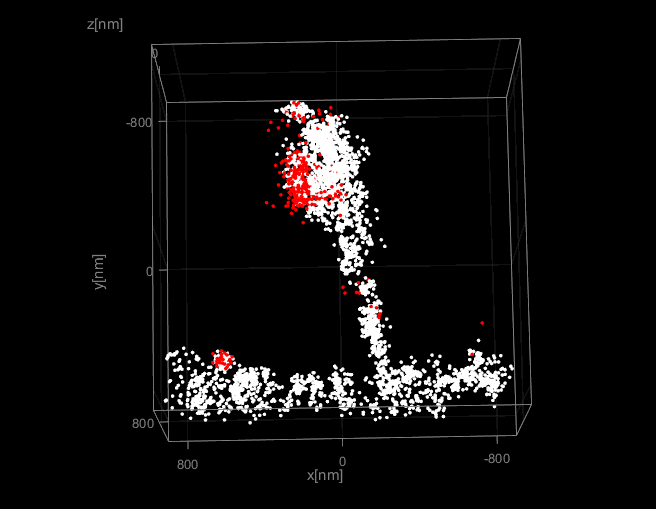

### MovieS2

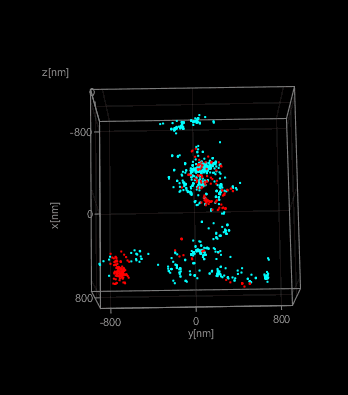

### MovieS3

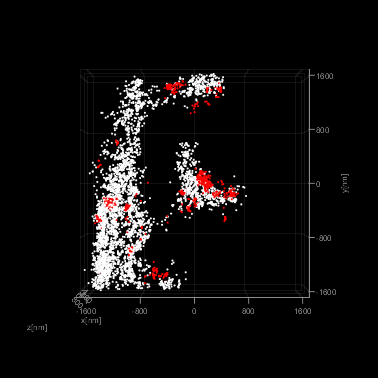
